# The chaperone Tsr2 regulates Rps26 release and reincorporation from mature ribosomes to enable a reversible, ribosome-mediated response to stress

**DOI:** 10.1101/2021.04.05.438496

**Authors:** Yoon-Mo Yang, Katrin Karbstein

## Abstract

Although ribosome assembly is quality controlled to maintain protein homeostasis, different ribosome populations have been described. How these form, especially under stress conditions that impact energy levels and stop the energy-intensive production of ribosomes, remains unknown. Here we demonstrate how a physiologically relevant ribosome population arises during high Na^+^ and pH stress via dissociation of Rps26 from fully assembled ribosomes to enable a translational response to these stresses. The chaperone Tsr2 releases Rps26 in the presence of high Na or pH *in vitro* and is required for Rps26 release *in vivo*. Moreover, Tsr2 stores free Rps26 and promotes re-incorporation of the protein, thereby repairing the subunit after the stress subsides. Our data implicate a residue in Rps26 involved in Diamond Blackfan Anemia in mediating the effects of Na^+^. These data demonstrate how different ribosome populations can arise rapidly, without major energy input, and without bypass of quality control mechanisms.

**Graphical Abstract:** 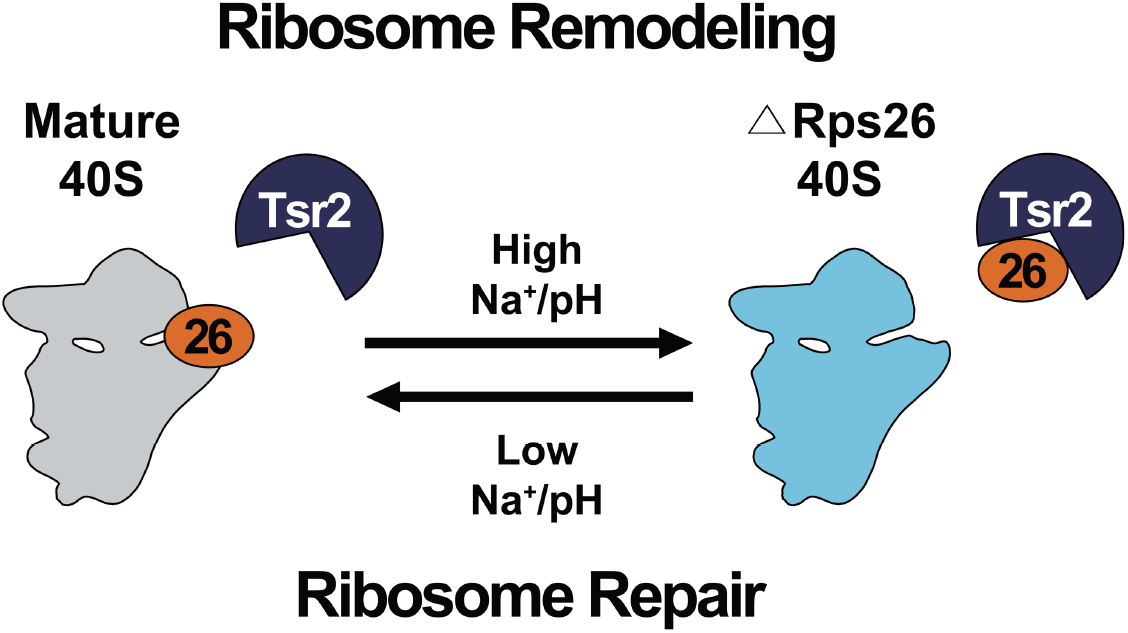

**Highlights:**

- Tsr2 releases Rps26 from mature ribosomes to remodel ribosome populations
- Tsr2 stores released Rps26 and allows for ribosome repair after stress
- Rps26 exploits a Mg binding site linked to Diamond Blackfan Anemia (DBA)
- Chaperone-mediated ribosome remodeling might be common for chaperoned RPs

## Introduction

Ribosomes synthesize proteins in all living organisms. Well over half of all transcription and translation events are dedicated to the synthesis of the 1000-2000 ribosomes/minute, which are required to *maintain* ribosome populations (Warner, 1999). This task is further complicated by the need to ensure that ribosomes are correctly assembled. For that purpose, cells have co-opted the translational machinery to establish a quality control pathway, which test-drives nascent ribosomes in a translation-like cycle (Lebaron et al., 2012; Strunk et al., 2012), and establishes check points to prevent immature or misassembled ribosomes from entering the translating pool (Ghalei et al., 2017; Huang et al., 2020; Parker et al., 2019). The importance of producing a homogeneous population of correctly assembled ribosomes is supported not just by these findings and the observation that mutations that bypass quality control occur in cancer cells (Huang et al., 2020; Parker et al., 2019; Sulima et al., 2014), but also from findings that cancer cells accumulate ribosomes with altered protein stoichiometry (Ajore et al., 2017; Guimaraes and Zavolan, 2016; Kulkarni et al., 2017; Vlachos, 2017). Moreover, insufficiency of ribosomal proteins can lead to the accumulation of ribosomes lacking these proteins (Cheng et al., 2019; Ferretti et al., 2017), and in human cells predisposes to cancer (Armistead and Triggs-Raine, 2014; Vlachos, 2017).

At the same time recent work has provided ample evidence for subpopulations of ribosomes that differ in composition (Emmott et al., 2019; Ferretti and Karbstein, 2019; Genuth and Barna, 2018a, b; Sauert et al., 2015). While many of these are associated with disease states and arise from haploinsufficiency of ribosomal proteins (Bolze et al., 2013; Fortier et al., 2015; Kondrashov et al., 2011), others are found in wild type cells (Ferretti et al., 2017; Loveland et al., 2016; Shi et al., 2017; Slavov et al., 2015). However, whether or not these have functional relevance, or are potentially even degradation intermediates is unclear in most cases (Ferretti and Karbstein, 2019). While ribosomes are costly to assemble, they are exceptionally stable and not turned over during the division cycle of most cells (Cole et al., 2009; LaRiviere et al., 2006; Warner, 1999). In the case of ribosomes deposited into the egg during oogenesis, these ribosomes must persist throughout the fertile life of the animal, which in the case of humans can be decades. Whether ribosomes become damaged over these extended time periods and then repaired remains unknown. Therefore, in addition to being functional, or represent nonfunctional degradation intermediates, ribosomes lacking individual ribosomal proteins could also represent intermediates of yet-to-be discovered repair pathways.

A ribosome sub-population with a known physiological role are Rps26-deficient ribosomes (Ferretti et al., 2017). Formed when yeast cells are exposed to high Na^+^ concentrations or high pH, they enable the translation of mRNAs containing an otherwise unfavorable guanosine residue at the −4 position of the Kozak sequence. This loss of preference towards the canonical −4A is explained by Rps26 interaction with this residue. About one quarter of all mRNAs in yeast are differentially bound by Rps26-deficient ribosomes, including those enabling the biological response to high Na^+^ and high pH (Ferretti et al., 2017). How these Rps26 deficient ribosomes are produced in cells is not known. In particular, it is unclear if they are newly made, and evading quality control mechanisms to ensure all proteins are incorporated, or if instead they are produced by release of Rps26 from pre-existing completely assembled ribosomes.

Using these physiological Rps26-deficient ribosomes as a case study, we report here how mature ribosomes can be remodeled under stress conditions and repaired once the stress subsides. Pulse-chase experiments demonstrate that Rps26-deficient ribosomes are formed by release of Rps26 from pre-existing, fully matured ribosomes, rather than by bypassing Rps26 incorporation into newly-made ribosomes. Furthermore, we show that Rps26 is re-incorporated once the stress is removed, demonstrating active repair of these ribosomes. Our experiments suggest that binding of metal ions and protons to specific sites at the Rps26-RNA interface allows for direct sensing of Na^+^ and H^+^ concentrations, and implicate Asp33, a residue mutated in Diamond Blackfan Anemia (DBA) in mediating salt-dependent effects. *In vivo* and *in vitro* experiments show that Rps26 release is enabled by the chaperone Tsr2, which has been previously suggested to deliver Rps26 to ribosomes (Schutz et al., 2014; Schutz et al., 2018). We show that Tsr2 promotes Rps26 dissociation and stores the released protein for reincorporation. This observation not only supports the importance of Tsr2 in adjusting the proper ratio of Rps26-containing and deficient ribosomes, but might also explain the conflicting observations of Tsr2 expression changes in cancer cells.

## Results

### Rps26-deficient ribosomes are generated from pre-existing mature ribosomes under stress

We have previously shown that Rps26-deficient ribosomes accumulate in yeast cells exposed to high Na^+^ concentrations or pH (Ferretti et al., 2017). What is unknown is whether these Rps26-deficient ribosomes arise by release of Rps26 from pre-existing mature ribosomes, or by omission of Rps26 during assembly of new ribosomes (**Figure 1A**). To distinguish between these pathways, we designed a pulse-chase experiment to differentiate between pre-made and newly-made ribosomes (**Figure 1A**). In this experiment, pre-existing ribosomes are marked with TAP-tagged Rps3 whose expression relies on the doxycycline (dox)-repressible TET promoter. Untagged Rps3 is under galactose-inducible/glucose repressible control. Cells were initially grown in glucose media, such that all pre-existing ribosomes will be marked with Rps3-TAP. In mid log phase, cells were switched to galactose media containing dox, such that newly-made ribosomes contain untagged Rps3. Control experiments confirm the rapid upregulation of Rps3 and downregulation of Rps3-TAP mRNAs under these conditions (**Figure S1**). NaCl was added when the cells are switched to galactose/dox media. Thus, ribosomes made before the stress (pre-existing ribosomes) are marked with Rps3-TAP, while newly-made ribosomes are untagged. These two ribosome populations were separated using IgG beads, and the relative levels of Rps26 in the pre-existing, TAP-tagged ribosomes (elution) and the newly-made ribosomes (flow-through) were measured by Western blotting in comparison with three other ribosomal proteins (Rps3, Rps8 and Rps10) as previously described (Ferretti et al., 2017). Importantly, Rps26 levels in pre-existing ribosomes (elution) were decreased approximately 50% (**Figure 1B-C**). This observation demonstrates that the Rps26-deficient ribosomes arise from release of Rps26 from pre-existing ribosomes when yeast are subject to high Na^+^ stress. This is consistent with our previous observation that in high Na^+^, Rps26 mRNA levels were downregulated as much but not more than other tested mRNAs (Ferretti et al., 2017), and the finding that stress conditions block ribosome assembly (Gasch et al., 2000; Warner, 1999), which would also block the de novo formation of Rps26-deficient ribosomes.

**Figure 1:**
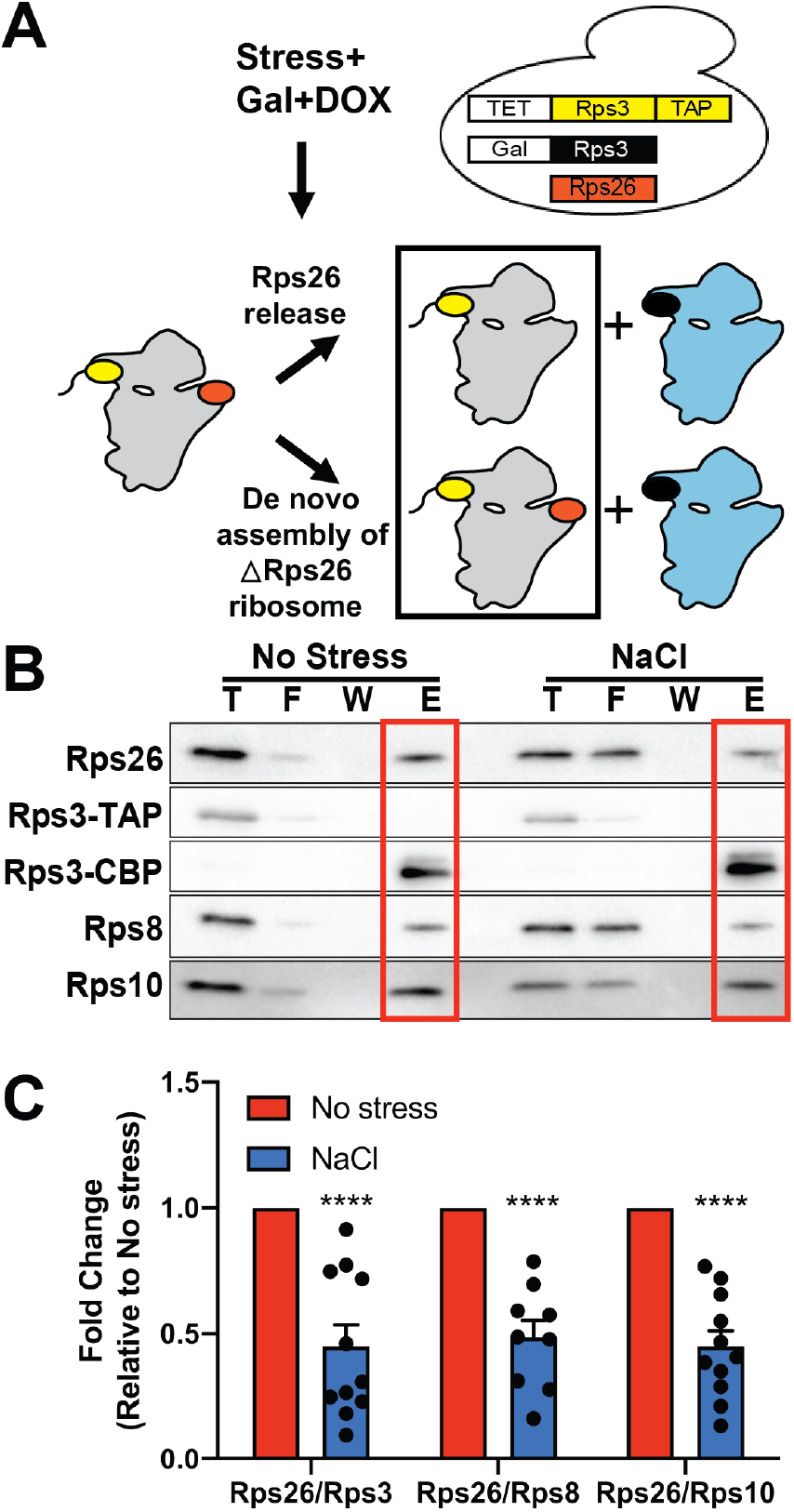
Rps26-deficient ribosomes arise from pre-existing 40S subunits. (A) Pulse-chase experiments to separate pre-existing ribosomes (grey) from newly-made ribosomes (blue) rely on a yeast strain with Rps3-TAP produced from a TET-repressible promoter (yellow), and Rps3 (black) produced from a galactose-inducible/glucose-repressible promoter. Rps26 is shown in red. By shifting this strain from glucose to galactose/dox, pre-existing ribosomes are marked with the Rps3-TAP affinity purification handle, and will be in the TAP-elution, as indicated by the black box. (B) Western blot of pre-existing (Rps3-TAP) ribosomes isolated by affinity purification from cells treated or not treated with 1M NaCl prior to lysis. T: total lysate; F: flowthrough; W: wash; E: elution. Note that elution from the IgG beads by TEV protease converts Rps3-TAP (Rps3-CBP-proteinA) to Rps3-CBP (Rps3-calmodulin-binding protein). (C) Quantification of data in panel (B). Data are averaged from 8 biological replicates with 1-2 technical replicates each. Error bars represent the SEM, and significance was determined using an unpaired t-test. ****, P < 0.0001.

### Tsr2 dissociates Rps26 from mature ribosomes in vitro

The data above demonstrate that Rps26 is released from pre-existing mature ribosomes to yield Rps26-deficient ribosomes under high salt stress. This led us to next ask how Rps26 dissociation occurs within cells. Tsr2 is a chaperone for Rps26, which stabilizes the protein outside of the ribosome (Peng et al., 2003; Schutz et al., 2014; Schutz et al., 2018). A role for Tsr2 in Rps26 incorporation into 40S subunits has also been suggested (Schutz et al., 2014; Schutz et al., 2018). Given Tsr2’s ability to bind and stabilize Rps26, we wondered if it could also release Rps26 from mature 40S. To test this hypothesis, we developed an *in vitro* release assay. After incubation of purified 40S subunits with purified recombinant Tsr2 in varying salt concentrations, the samples were loaded onto a sucrose cushion and spun at high speed to pellet ribosomes (and its bound Rps26), thereby separating it from released Tsr2-bound Rps26, which will remain in the supernatant. The data in **Figure 2A-B** demonstrate that at increased concentrations of KOAc, Tsr2 releases Rps26 from mature 40S subunits. Release from the 40S subunit was specific for Rps26, as other ribosomal proteins were observed in the pellet fraction (**Figure 2A, S2A**-**B**). Finally, even though Tsr2-independent release of Rps26 was observed at 1M KOAc (**Figure 2C-D**), the results here show that at moderate KOAc concentrations the release of Rps26 is promoted by Tsr2.

**Figure 2:**
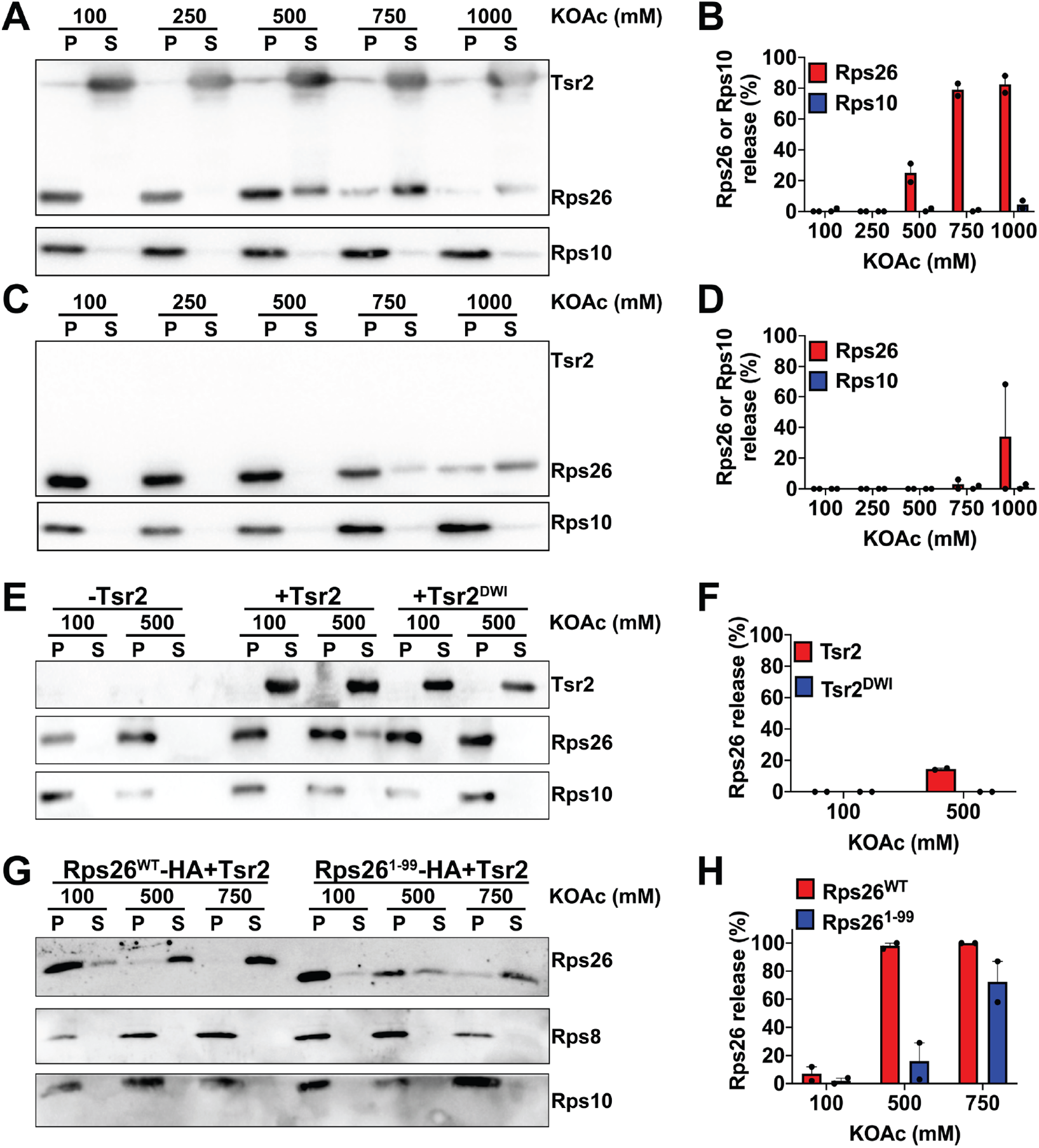
Tsr2 promotes Rps26 release from mature 40S subunits in the presence of high salt *in vitro*. Western blot analysis of pelleted (P) ribosomes and released proteins in the supernatant (S). Mature 40S subunits purified from yeast were incubated with (A) or without (C) recombinant Tsr2 under different salt concentrations. (B and D) Released Rps26 or Rps10 was quantified from panels (A) and (C) with two independent replicates for each condition. Error bars represent the SEM. (E) Rps26 release assay with the Rps26-interaction-deficient Tsr2^DWI^ mutant. (F) Quantification of Rps26 in panel (E) with two independent replicates for each condition. Error bars represent the SEM. (G) Rps26 release assay with the Tsr2-interaction-deficient Rps26^1-99^ (lacking residues 100-119) truncation mutant. (H) Quantification of released Rps26 in panel (G) with two independent replicates for each condition. Error bars represent the SEM.

To verify that the physical interaction between Tsr2 and Rps26 is essential for Rps26 dissociation and demonstrate a specific interaction between Tsr2 and Rps26, we wanted to confirm that Rps26 release required the characterized binding interface between Rps26 and Tsr2. Previous work has shown that the Tsr2^DWI^ mutant (D64W65I66A) impairs binding to Rps26 (Schutz et al., 2018). As expected, Tsr2^DWI^ was unable to dissociate Rps26 from 40S subunits at high salt concentrations (**Figure 2E-F**).

Similarly, the C-terminal tail of Rps26 is required for binding to Tsr2 (Schutz et al., 2018). If Tsr2 released Rps26 from ribosomes by binding this part of the protein, we would expect that truncated Rps26^1-99^ is resistant to release from 40S subunit under increased level of KOAc. Indeed, the release of Rps26 was more efficient from 40S subunits containing full length Rps26^WT^ compared to those containing Rps26^1-99^ (**Figure 2G-H**). Thus, these data demonstrate that the Tsr2-dependent release of Rps26 from 40S subunits requires binding of Tsr2 to Rps26 via its previously characterized binding interface.

### Tsr2 remodels mature ribosomes to generate Rps26-deficient ribosomes under stress

Above, we have shown that Tsr2 can release Rps26 from mature ribosomes in the presence of high salt *in vitro*. We next tested if Tsr2 was also involved in Rps26 release from ribosomes when yeast cells are stressed *in vivo*. If so, then we predict that the released Rps26 from pre-existing ribosomes will be bound to Tsr2 under stress. To confirm this hypothesis, we redesigned the pulse-chase experiment by following pre-existing Rps26-HA bound to Tsr2-TAP. In this experiment, the expression of HA-tagged Rps26 is regulated by a galactose-inducible/glucose-repressible promoter (in the background of constitutive untagged Rps26). By switching the media to YPD when salt stress is applied, we ensure that only pre-existing ribosomes contain Rps26-HA (**Figure 3A**). If Tsr2 binds Rps26-HA released from pre-existing ribosomes, then we expect that more pre-existing Rps26-HA is bound to Tsr2 in stress-treated than untreated cells. The data in **Figure 3B-C** show that indeed more of the pre-existing Rps26-HA is bound to Tsr2 in stress-treated than untreated cells. The same is true for untagged Rps26, as expected as this is a mixture of new and pre-existing Rps26. Thus, these data show that after its release from 40S subunits, Rps26 is bound to Tsr2.

**Figure 3:**
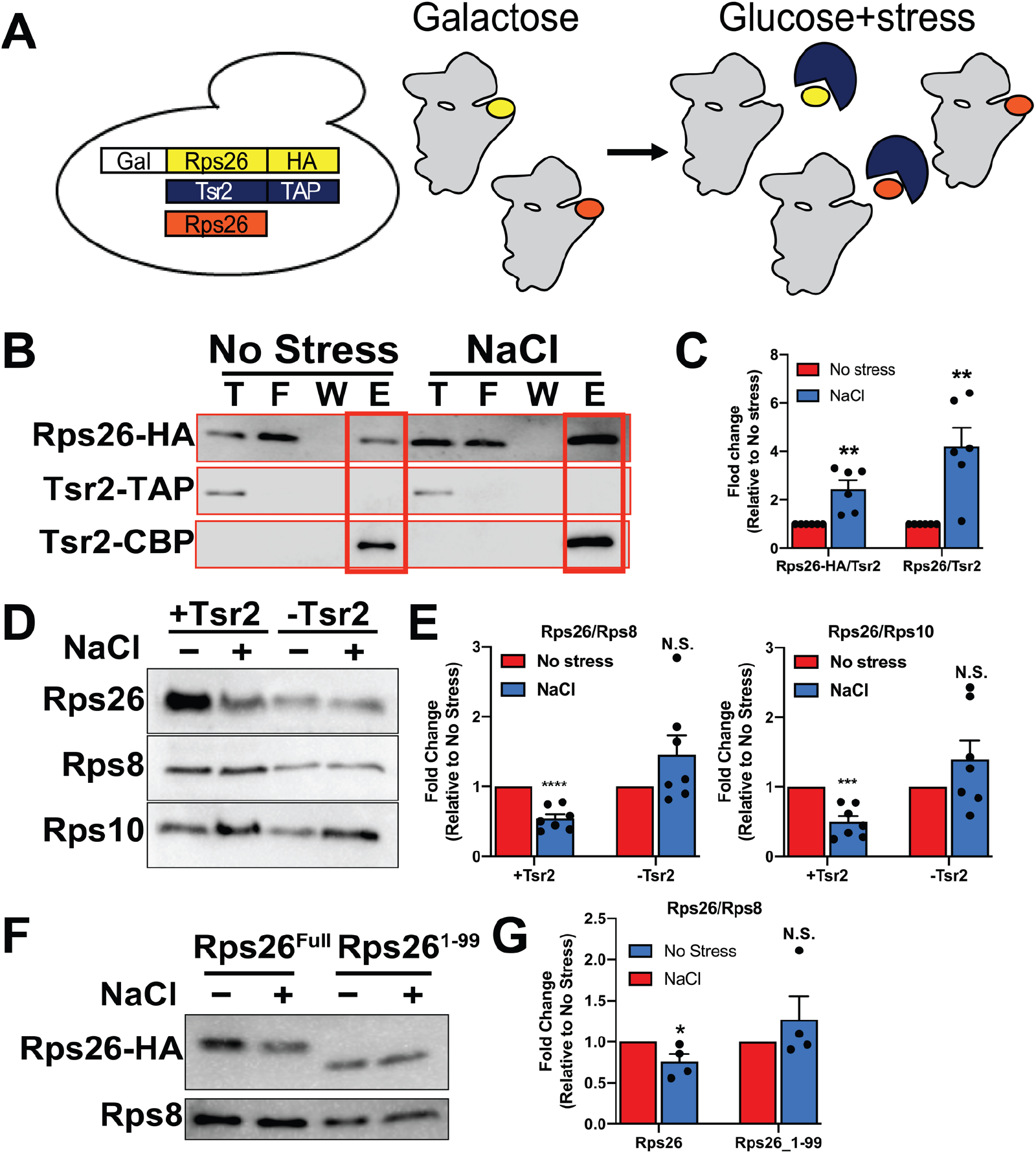
Tsr2 binds and dissociates Rps26 from pre-existing ribosomes under salt stress *in vivo*. (A) Pulse-chase experiments to follow pre-existing Rps26-HA (yellow) rely on a yeast strain where Rps26-HA is produced from a galactose-inducible/glucose-repressible promoter. Tsr2 (blue) is captured with the TAP-affinity tag, and untagged Rps26 is in red. (B) Western blot of pre-existing Rps26-HA co-isolated with TAP-tagged Tsr2 from cells that were or were not treated with 1M NaCl. (C) Quantification of data in panel (B). Data are averages from 6 biological replicates. Error bars represent the SEM. Significance was determined using an unpaired t-test. **, P < 0.01. (D) Rps26 concentrations in cell lysates from wild type or Tsr2-depleted cells. BY4741 (+Tsr2) or Gal::Tsr2 (−Tsr2) cells grown in glucose media were or were not treated with 1M NaCl. Rps8, Rps10 and Rps26 levels in cell lysates were determined by Western blot. (E) Quantification of data in panel (D). Data are averages from 7 biological replicates. Error bars represent the SEM. Significance was determined using an unpaired t-test. P < 0.001; ****, P < 0.0001. (F) Rps26 depletion in ribosomes from cells with full-length or truncated Rps26. Cells containing full-length or truncated, Tsr2-binding deficient Rps26^1-99^ were or were not treated with 1M NaCl. The concentrations of Rps8 and Rps26 in total cell lysates were measured by western blot. (G) Quantification of data in panel (F). Data are averages from 4 biological replicates. Error bars represent the SEM. Significance was determined using an unpaired t-test. *, P<0.05.

To demonstrate that Tsr2 was involved in the release of Rps26, and not just storing Rps26 that was released by other means, we compared the amount of Rps26 in stress-treated and untreated cells when Tsr2 is present or absent in those cells. These experiments demonstrate that Tsr2 is required for the reduced levels of Rps26 after Na^+^ addition, as no change or even an increase in Rps26 is observed in the absence of Tsr2 (**Figure 3D-E)**.

Moreover, as for the *in vitro* experiments, the C-terminal tail of Rps26, which binds Tsr2, is required for the change in Rps26 levels with NaCl-stress, as Rps26 levels remain constant in Rps26^1-99^ cells, while they are reduced in cells containing wild type Rps26 (**Figure 3F-G)**.

Thus, Tsr2 and its previously characterized binding interface on Rps26 are required to generate Rps26-deficient ribosomes *in vitro* and *in vivo*. Together with the observation that Tsr2 can release Rps26 *in vitro* and that released Rps26 is bound to Tsr2 *in vivo*, these data show that Tsr2 is the release factor for Rps26 *in vivo*.

### Rps26-deficient ribosomes are repaired to mature ribosomes after stress

Above we have shown that during high salt or pH stress Rps26 is released from pre-existing ribosomes by Tsr2, to which it remains bound. In addition, previous work suggested that Tsr2 is involved in the incorporation of Rps26 into ribosomes. We therefore wondered if the Tsr2-dependent release of Rps26 was reversible, such that Rps26 could be reincorporated from the Tsr2-bound complex into ribosomes once stress was removed. This would spare ribosomes whose assembly requires extensive resources from destruction, while also helping cells to quickly return to the normal translational program.

To test if Rps26-deficient ribosomes can re-incorporate Rps26 *in vitro*, we took advantage of our release assay. We first generated 40S subunits lacking Rps26 by incubating 40S subunits with Tsr2 in high salt, and separating them from released Rps26•Tsr2 (**Figure 4A**, Rps26 dissociation). We then added recombinant Tsr2·Rps26 to the Rps26-deficient ribosomes in low (100 mM) or high (750 mM) K^+^. Rps26 was re-incorporated only when the K^+^ concentration was lowered, showing that Rps26 can assemble into Rps26-deficient ribosomes from the Tsr2•Rps26 complex (**Figure 4A**). Moreover, we note that regeneration is complete, as half of Rps26 was observed in the pellet after addition of a two-fold excess in the Tsr2·Rps26 complex.

**Figure 4:**
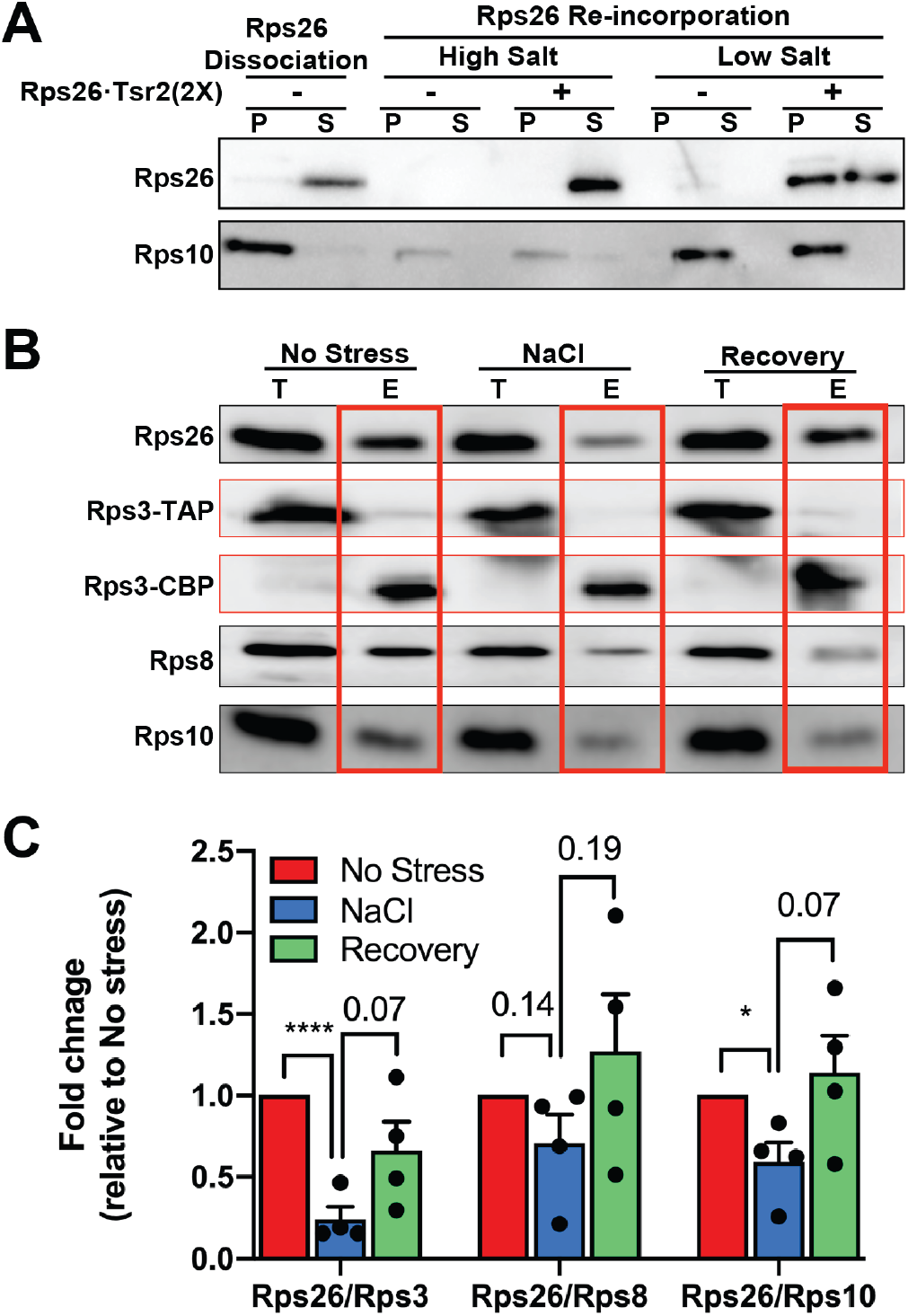
Repair of Rps26-deficient ribosomes after stress. (A) Rps26-cosedimentation with ribosomes after incubation of Rps26-deficient ribosomes with recombinant Tsr2•Rps26 *in vitro*. Western blot analysis of ultracentrifugation pelleting experiments. Rps26-deficient ribosomes (Rps26 Dissociation) were generated by Tsr2 addition in 750 mM KOAc and separated from free Rps26•Tsr2 by ultracentrifugation. Rps26-deficient ribosomes were incubated with purified recombinant Rps26•Tsr2 in low (100 mM KOAc) or high (750 mM KOAc) salt buffers, and Rps26-binding was assessed after pelleting of the ribosomes. S: supernatant; P: pellet. (B) Rps26 is reinserted into ribosomes *in vivo*. To monitor the repair of Rps26-deficient ribosomes by re-insertion of Rps26, the same pulse-chase experiment as in Figure 1 was performed, except NaCl stress was relieved by dilution into fresh media. T: total lysate; E: elution. (C) Quantification of data in panel (B). Data are averages from 4 biological replicates. Error bars represent the SEM, and significance was determined using an unpaired t-test. *, P < 0.05; ****, P < 0.0001.

To test if Rps26-deficient ribosomes can be repaired by re-incorporation of Rps26 from Tsr2•Rps26 *in vivo*, we extended the pulse-chase experiment in Figure 1. As before, we first generated Rps26-deficient 40S subunits by addition of high salt to yeast cells (**Figure 4B-C**, NaCl), before removing the stress for 1h. After this recovery, Rps26 levels in pre-existing ribosomes also recovered (**Figure 4B-C**, recovery), showing that Rps26 can be reincorporated into pre-existing ribosomes from which it had been previously released.

Together, the results here suggest that Tsr2 releases Rps26 from ribosomes during high salt to form a Tsr2•Rps26 complex. Moreover, the data also indicate that these ribosomes can then be repaired, by reincorporation of Rps26 from the Tsr2•Rps26 complex.

### Ribosomes directly sense increased Na^+^/pH *in vitro*

Above we have shown that addition of 1M NaCl leads to the release of Rps26 from ribosomes to Tsr2 *in vivo*, which can be reversed when the salt stress is removed. We next wondered if the function of Tsr2 in either releasing or incorporating Rps26 into 40S subunits was simply a function of the salt concentration, or pH, or perhaps required additional signaling cascades, resulting in posttranslational modifications such as phosphorylation or ubiquitination. To search for evidence of large posttranslational modifications, such as ubiquitination, *in vivo* we used Western blotting after salt exposure. However, these data do not provide any evidence for ubiquitination of either Rps26 or Tsr2 when yeast cells were treated with NaCl stress (**Figure S3A**). Moreover, no increase in phosphorylation of Rps26 or Tsr2 was observed when phosphorylation of the total proteome was analyzed after NaCl treatment (MacGilvray et al., 2018), and we were unable to detect phosphorylation of Rps26 or Tsr2 using phosphoserine antibodies, (**Figure S3B-C**). While these (or any other) data cannot rule out a distinct posttranslational modification, they led us to consider whether the ribosome could directly sense an increase in the salt concentration or pH, becoming susceptible to Tsr2-mediated extraction even at physiological Na^+^ concentrations. This model would be consistent with the previous observation that the Tsr2•Rps26 complex was strengthened at high salt concentrations (Schutz et al., 2018), while the Rps26•40S interaction is weakened **(Figure 2C)**.

While the data above show that Tsr2-mediated Rps26 release required about 500 mM KOAc, the concentrations of Na^+^ and K^+^, the two most abundant cations in yeast cells, are ~20mM and ~200mM, respectively (Larsson et al., 1998). Addition of 1M NaCl to the media leads to an increase in the intracellular Na^+^ concentration to ~150 mM, while the total concentration of Na^+^ and K^+^ is maintained at ~300 mM by decreasing the intracellular level of K^+^ to ~150 mM (Ferrando et al., 1995; Larsson et al., 1998). We therefore wondered if Na^+^ and K^+^ had different concentration dependences for Rps26 release, reflecting different affinities, as expected from an organized binding site that has a regulatory function.

To test this hypothesis, we compared Na^+^ and K^+^ effects on Rps26 release. The data show that Na^+^ concentrations that lead to Rps26 dissociation are much lower than K^+^ concentrations (**Figure 5A-B**), and within the range of physiological intracellular concentrations under high salt stress. This result shows that Tsr2 can dissociate Rps26 at the Na^+^ concentrations found within salt-treated yeast cells.

**Figure 5:**
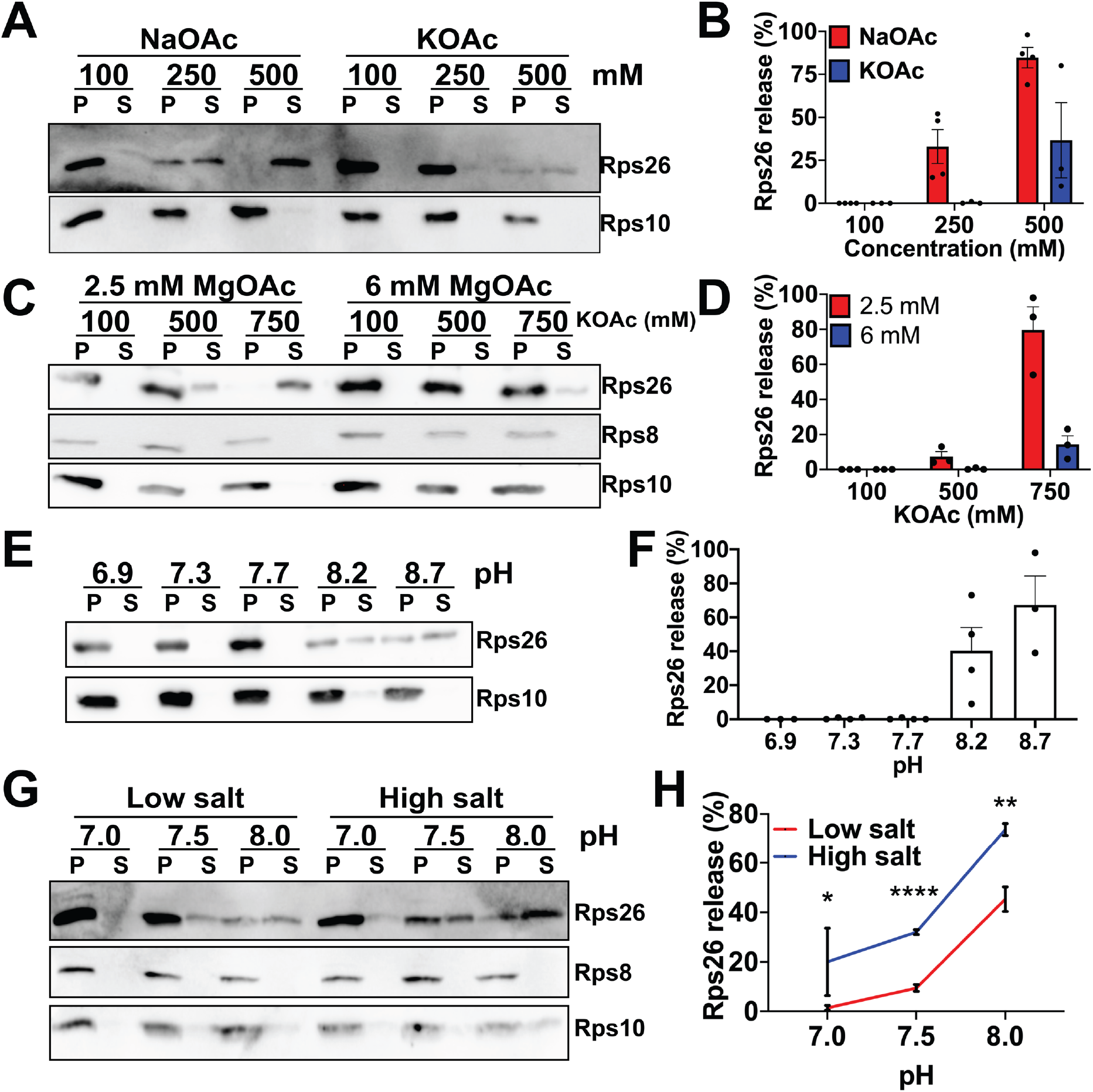
Ribosomes directly sense ion and protein concentrations *in vitro*. (A) Western blot analysis of pelleted (P) ribosomes and released proteins in the supernatant (S). Mature 40S subunits purified from yeast were incubated with recombinant Tsr2 at different concentrations of NaOAc or KOAc. (B) Quantification of Rps26 in panel (A) with 3-4 independent replicates for each condition. Error bars represent the SEM. (C) Effect of Mg^+^ in the release of Rps26. Mature 40S subunits were incubated with recombinant Tsr2 in either 2.5 mM or 6 mM MgOAc at different KOAc concentrations. (D) Quantification of Rps26 in panel (C) with three independent replicates for each condition. Error bars represent the SEM. (E) Release assay in 20 mM Bis-Tris propane at various pH values, 2.5 mM MgOAc and 100 mM KOAc. (F) Quantification of Rps26 in (E) with 3-4 independent replicates for each condition. Error bars represent the SEM. (G) Release of Rps26 under physiological conditions. Mature 40S subunits were incubated with recombinant Tsr2 in either low physiological salt conditions (20 mM NaCl, 200 mM KCl, 2.5 mM MgOAc) or high physiological salt concentrations (150 mM NaCl, 150 mM KCl, 2.5 mM MgOAc) with different pH values adjusted by Bis-Tris propane. (H) Quantification of results in panel (G) with 3-9 independent replicates for each condition. Error bars represent the SEM. Significance was determined using an unpaired t-test. *, P < 0.05; **, P < 0.01; ****, P < 0.0001.

To further dissect how K^+^ or Na^+^ affected Rps26 binding, we next tested the model that K^+^ or Na^+^ evict a Mg^2+^ ion stabilizing Rps26 binding. Thus, we repeated the release assay at higher Mg^2+^ concentrations. Increasing the Mg^2+^ concentration from 2.5 to 6 mM largely blocks the Na^+^-dependent release of Rps26 (**Figure 5C-D**), supporting the model that increased intracellular salt concentrations lead to Rps26 release from 40S subunits via competition with a Mg^2+^ ion critical for Rps26 binding.

In addition to NaCl stress, alkaline pH stress also promoted the formation of Rps26-deficient ribosomes in yeast cells (Ferretti et al., 2017). Therefore, we further tested whether pH changes could also trigger Tsr2-dependent Rps26 release *in vitro*. Increasing the pH of the growth media to pH 8.2, will increase the intracellular pH to ≥7.5 ((Orij et al., 2009); as the authors note the measured pH of 7.5 is likely an underestimate). We therefore varied the pH in release experiments between pH 6.9 and pH 8.7. Increasing the pH leads to Tsr2-dependent Rps26 dissociation from mature ribosomes (**Figure 5E-F, Figure S2E)**. These data strongly suggest that changes in pH, like changes in Na^+^ concentration, are also directly recognized by the ribosome leading to Rps26 dissociation.

Finally, we asked whether changes in pH and salt were acting via the same residues, or different residues, which would make these changes in the cellular environment additive. To answer this question, we tested whether Rps26 release was affected by pH changes on top of ion changes, or whether at high pH, salt would no longer matter. As shown in **Figure 5G-H**, changes in pH and salt are clearly additive, suggesting that they act via distinct residues at the 40S•Rps26 interface. Moreover, these observations also suggest that the response to these physiological changes is synergistic, such that lower salt is required when the pH is also elevated. Finally, the data show that under physiological low salt conditions the elevated pH 7.5 observed in pH stress cells can effect Rps26 release, while at pH of 7.0 the physiological high salt conditions lead to Rps26 release (**Figure 5G-H**).

## Discussion

### Tsr2-dependent remodeling and repair of ribosomes by reversible release of Rps26

We have previously shown that yeast cells exposed to high Na^+^ or high pH stress accumulate ribosomes lacking Rps26 (Ferretti et al., 2017). These enable translation of mRNAs with an otherwise unfavorable G at position −4 in the Kozak sequence, thereby promoting a distinct translational program that supports the response to these stresses (Ferretti et al., 2017). Here we address how these Rps26-deficient ribosomes are formed (**Figure 6A**).

**Figure 6.**
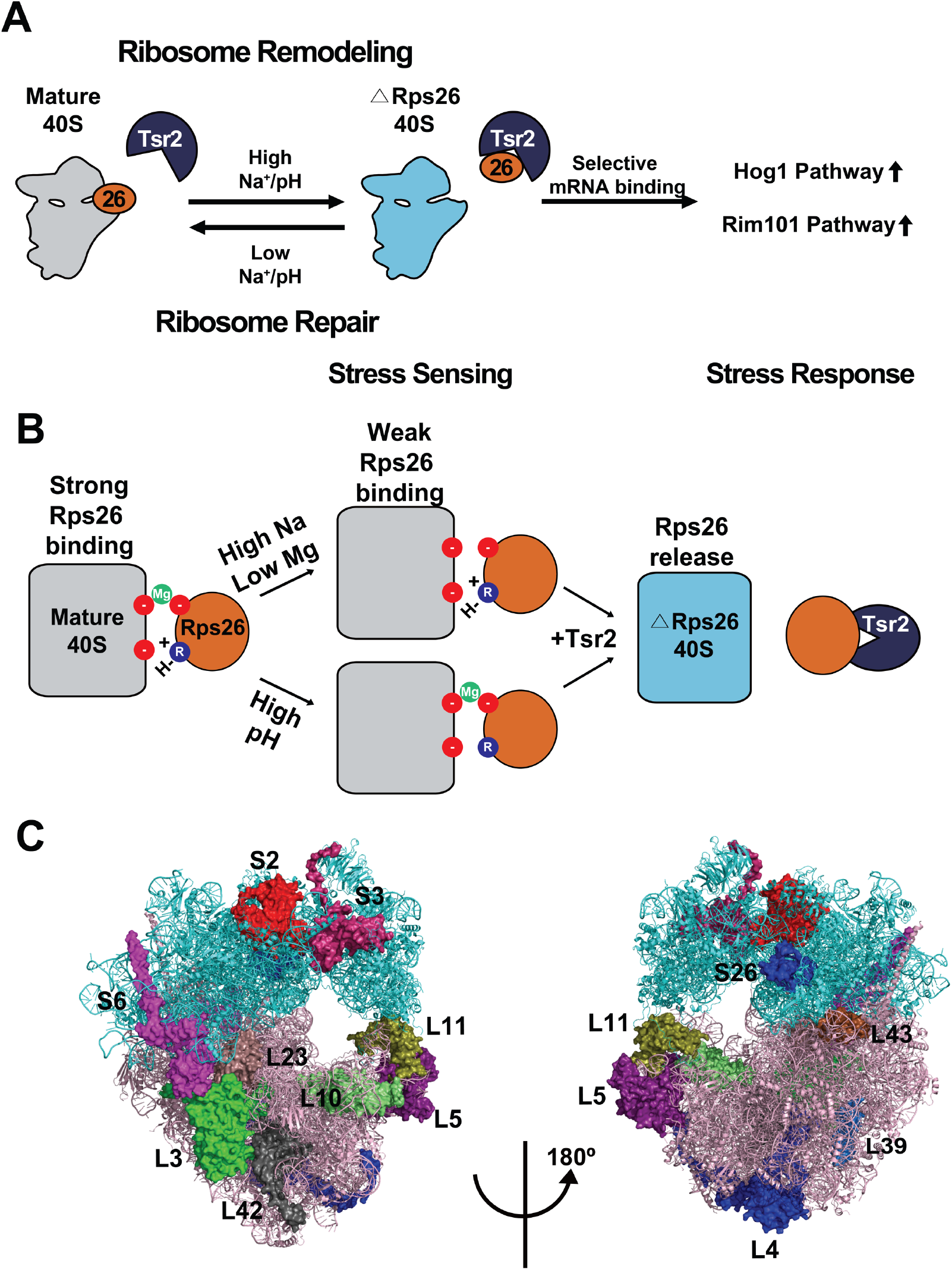
Model for the Tsr2-mediated remodeling and repair of Rps26-deficient ribosomes. (A) High intracellular Na^+^ concentrations or pH leads to ribosome remodeling by Tsr2-dependent dissociation of Rps26 from 40S ribosomes, producing Rps26-deficient ribosomes that display altered mRNA preference and enable translation of mRNAs from the Hog1 and Rim101 salt and pH response pathways (Ferretti et al., 2017). Removal of the stress allows for ribosome repair by allowing reincorporation of Rps26 from the Rps26•Tsr2 complex. (B) Rps26 binding to 40S ribosomes is Mg^2+^ and H^+^ dependent, and involves two distinct sites. (C) Ribosomal proteins with specialized chaperones cluster in functionally important regions. PDB 4V88. The small and large subunits are shown in cyan and pink, respectively. Chaperoned proteins are highlighted in space fill.

Our data show that changes in the salt or pH affect the 40S subunit directly, weakening the binding of Rps26 such that it can be released by Tsr2, a small acidic protein that can form a stable complex with Rps26, thereby stabilizing Rps26 outside of the ribosome. Thus, Rps26-deficient ribosomes are formed by Tsr2-dependent release of Rps26 from pre-existing ribosomes. Mutations in the Tsr2/Rps26 interface impair Rps26 release *in vivo* and *in vitro*.

Moreover, we also demonstrate that Rps26 can be incorporated from the Tsr2•Rps26 complex into Rps26-deficient ribosomes *in vivo* and *in vitro*, thereby providing direct evidence for a role of this chaperone in incorporation of Rps26. Such a role was previously suggested based on the observation that Tsr2 stabilizes Rps26 outside of the ribosome (Schutz et al., 2014), and that the growth defect from Tsr2 deletion can be suppressed by Rps26 overexpression (Schutz et al., 2018).

Together, these results suggest that under stress Tsr2 remodels 40S ribosomal subunits to release Rps26, which is stored in the Tsr2•Rps26 complex, from which it can be reincorporated to repair the subunits after the stress subsides (**Figure 6A**). To our knowledge, this is the first instance of active remodeling and repair of ribosomes, prompting us to speculate that ribosome remodeling could be a more common way to rapidly generate different ribosome populations, and that repair of ribosomes damaged by age (*e*.*g*. in oocytes) or by oxidative stress could be observable in other instances.

### Ribosomes as sensors for fluctuations in intracellular salt, Mg^2+^ and pH

The results in Figure 5 show that the mature 40S subunit itself can detect variations in Mg^2+^, Na^+^ or H^+^ concentrations, triggering the Tsr2-enabled dissociation of Rps26. The Rps26-deficient ribosomes then function by enabling the translation of mRNAs with an otherwise disfavored −4G mutation in the Kozak sequence, which changes protein homeostasis (Ferretti et al., 2017). Thus, the ribosome is both a sensor for physiological changes in intracellular salt, Mg and pH, as well as a mediator for responding to these changes (**Figure 6A**).

How do ribosomes sense the differences of salt and pH? Release assays at different Mg^2+^ concentrations strongly suggest that the release of Rps26 by Na^+^ occurs via competition with a Mg^2+^ ion that stabilizes Rps26 binding, either directly by bridging the RNA and protein, or by stabilizing the RNA structure (**Figure 6C**). Crystal structures identify multiple Mg^2+^ ions near the Rps26 binding site (Ben-Shem et al., 2011), including one bound to Asp33, an essential residue conserved in human Rps26 (**Figure S4A-B**; at 3Å resolution it is impossible to distinguish Mg^2+^ from K^+^ ions; however, the crystal well contains fairly high concentrations of Mg^2+^ (3.3-10 mM) and low concentrations of K^+^ (95 mM)). This Mg^2+^ also contacts the rRNA backbone and could thus be a candidate ligand for mediating the salt-dependent effect. Intriguingly, the D33N mutation leads to Diamond Blackfan Anemia (Doherty et al., 2010), but does not affect the interaction with Tsr2 (Schutz et al., 2014). We thus hypothesized that the D33N mutation weakens Rps26 binding via loss of the Mg^2+^ binding site. Indeed, purification of 40S subunits from cells containing Rps26_D33N shows that these have lost nearly all Rps26 (**Figure S4C)**, demonstrating the importance of the metal ion at that position for Rps26 binding. Because these ribosomes entirely lack Rps26, we could not directly test if Rps26-release displayed a different response to high salt. Nonetheless, we note that there are two additional ions within less than 7Å to the D33 bound ion (**Figure S4A**). It is conceivable that the differential occupation of these sites by Na^+^, K^+^ and Mg^2+^ is responsible for the observed salt effects on Rps26 release.

The data also demonstrate that salt and pH act independently of each other, suggesting that the pH-sensitive site is distinct from the salt-dependent site (**Figure 6B**). Consistently, our data indicate a pKa value above 8 for the ionizable group(s), implicating side-chains such as lysines and histidines, which are not typically metal ligands.

### Remodeling pre-existing ribosomes might be advantageous under stress

Transcriptional responses to remodel the proteome under stress are well characterized and occur rapidly (Gasch et al., 2000). Thus, the advantages of changing the ribosome population are not immediately clear but could include the ability to produce different branches of a stress response by overlaying it onto changes in the mRNA population (Brar and Weissman, 2015; Cheng et al., 2019; Ferretti and Karbstein, 2019), as well as the potential to preferentially boost production of the stress proteome while producing moderate, instead of very high amounts of the corresponding mRNAs. Nonetheless, the potential disadvantage, the need to turn over an entire ribosome population at great energetic cost, is clear. This is especially puzzling, because energy supplies are often limited under cellular stress and because stress generally stops ribosome transcription and assembly (Gasch et al., 2000; Warner, 1999). Moreover, the turnover of ribosomes is also expected to be much slower than the turnover of mRNAs, thereby rendering such a response relatively slow (Cole et al., 2009; LaRiviere et al., 2006). Here we show how these disadvantages can be neutralized by remodeling pre-existing ribosomes to rapidly produce distinct ribosome populations without any additional energy input. Notably, ribosome remodeling also solves an additional potential problem, the existence of quality control pathways in place to ensure that ribosomes are correctly assembled (Ghalei et al., 2017; Huang et al., 2020; Lebaron et al., 2012; Parker et al., 2019; Strunk et al., 2012). Presumably, such mechanisms limit the accumulation not just of ribosomes lacking head components (Huang et al., 2020), but also those lacking Rps26. By generating Rps26-deficient ribosomes from fully and correctly assembled subunits, such quality control mechanisms do not need to be circumvented, which could open the cells to the uncontrolled production of Rps26-deficient ribosomes, which are associated with DBA (Boria et al., 2010).

### Are there other examples of ribosome remodeling and repair?

The work herein shows how at high Na^+^ concentrations or high pH, Rps26 can be extracted by its chaperone Tsr2 to yield Rps26-free ribosomes. Moreover, these can also be reconstituted to Rps26-containing ribosomes. *In vivo* this occurs when yeast cells are exposed to high salt or pH. Similarly, ribosomes from bacteria in stationary growth phase contain bL31B and bL36B instead of bL31A and bL36A, respectively ((Lilleorg et al., 2019), **Table 1**). While it wasn’t shown that the difference involved exchange of the bL31 and bL36 proteins, rather than replacement of ribosomes, bL31A can be exchanged for bL31B at low pH *in vitro*, suggesting the same to occur *in vivo*. Similarly, pulse-chase experiments with radioactively labeled proteins show binding of labeled bL31, uL1/Rpl1, uL5/Rpl11, uL10/Rpp0, uL11/Rpl12, uL30/Rpl7, uS2/Rps0 and uS5/Rps2 as well as bL9, bL33 and bS21 into ribosomes under conditions where assembly does not occur ((Pulk et al., 2010; Robertson et al., 1977; Subramanian and van Duin, 1977); note that proteins with homologs in bacteria and eukaryotes are labeled with the uL/Rpl or uS/Rps nomenclature; proteins without eukaryotic homologs are labeled with the bL/bS nomenclature and those without bacterial homologs with the Rpl/Rps nomenclature). Consistently, N-labeled bS20, bS21 and bL33 are also detected in assembled ribosomes faster than expected in pulse-chase mass-spectrometry experiments, suggesting that they exchange into pre-existing ribosomes and not newly-made ribosomes (Chen et al., 2012). Of note, while bS21 and Rps26 share no discernable homology, they bind at the same location (**Figure S5A**). While it may be unlikely that all of these proteins are exchangeable, we note that for three of them, bS21, bL31 and bL33 multiple independent studies provide evidence for exchange as described above, bolstering this evidence (**Table 1**). Nonetheless, evidence for extraction, as opposed to spontaneous dissociation and equilibration, has not yet been provided.

**Table 1:**
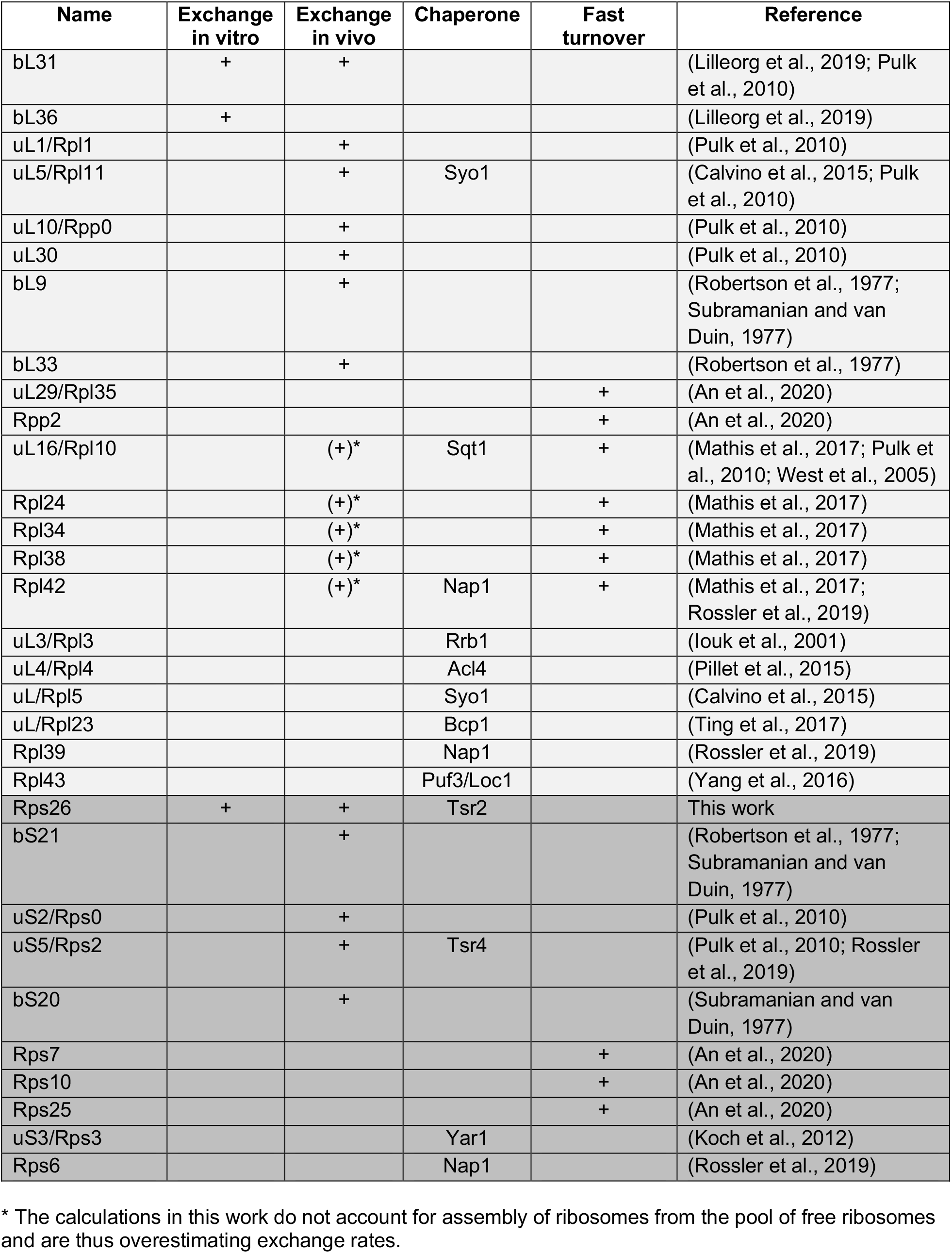
Ribosomal proteins that may be exchangeable

While exchange of ribosomal proteins has not been demonstrated in eukaryotes, in mammalian cells uL29/Rpl35, Rpp2, Rps7, and Rps10 as well as uL16/Rpl10, Rpl24, eL34/Rpl34, Rpl36A/Rpl42, and Rpl38 turn over faster than the rest of the ribosomal proteins (An et al., 2020; Mathis et al., 2017), indicating that they could be exchanged and then degraded from a free pool. Thus, exchange of ribosomal proteins between a free and a ribosome-bound pool appears possible for a sizeable subset of these proteins (**Table 1**).

Most ribosomal proteins are chaperoned by the translation-machinery associated general chaperones NAC and Ssb/RAC, which capture these proteins co-translationally and accompany them to the nascent ribosome (Koplin et al., 2010). This efficient system functions well for nearly all ribosomal proteins and prevents them from aggregating prior to their incorporation into ribosomes. Nonetheless, besides Tsr2, 11 more ribosomal protein-specific chaperones are known to date: Rps2:Tsr4, Rps3:Yar1, Rps6:Nap1, Rpl3:Rrb1, Rpl4:Acl4, Rpl5:Rpl11:Syo1, Rpl10:Sqt1, Rpl23:Bcp1, Rpl39:Nap1, Rpl42:Nap1 and Rpl43:Puf6:Loc1 ((Black et al., 2019; Calvino et al., 2015; Iouk et al., 2001; Koch et al., 2012; Pillet et al., 2015; Rossler et al., 2019; Ting et al., 2017; West et al., 2005; Yang et al., 2016), note that it is also possible that Nap1 forms a multimeric complex with Rps6, Rpl39 and/or Rpl42 at the same time), raising the question why this highly complex system has evolved in parallel to NAC/RAC. For some proteins, the chaperones function as nuclear import adaptors (Calvino et al., 2015; Iouk et al., 2001; Kressler et al., 2012; Mitterer et al., 2016; Schutz et al., 2014; Stelter et al., 2015), which could provide an advantage. However, NAC/RAC can enter the nucleus with all other ribosomal proteins, raising doubt about the importance of this contribution. We suggest that personalized chaperones allow not just for the individually regulated incorporation of ribosomal proteins, but also their extraction, as we have shown here for Tsr2-Rps26. Indeed, given the existence of checkpoints to ensure that all ribosomal proteins are incorporated into nascent subunits (Huang et al., 2020), making it difficult to leave out proteins, individually regulated extraction may be more relevant.

Most of the ribosomal proteins that have personalized chaperones (except Rpl4, Rpl42) are located on the surface of the ribosome, suggesting that they could be extracted by a chaperone akin to Rps26 (**Figure 6C**). Moreover, evidence of exchange exists for bacterial uL5/Rpl11 and uS5/Rps2 as well as mouse uL16/Rpl10 and Rpl42 (**Table 1**). Interestingly, the proteins with chaperones are all located in functionally important regions: three of the four small subunit proteins line the mRNA binding channel (Rps2, Rps3 and Rps26); two are located in the peptidyltransferase center (Rpl3, Rpl10), two (Rpl4, Rpl39) line the peptide exit tunnel and five proteins (Rps6, Rpl3, Rpl11, Rpl23 and Rpl43) are at the subunit interface (**Figure 6C**). Thus, their exchange or removal may affect ribosome function.

What rationale (other than repair of these functionally important centers) would there be for chaperone-mediated extraction of ribosomal proteins? One possibility is the modulation of ribosome function via turnover of posttranslationally modified ribosomal proteins (RPs): *e*.*g*. Rpl3 is methylated at His243, which supports translational fidelity (Webb et al., 2010). While histone (lysine) or DNA/RNA demethylases are known, no such enzyme has been described for histidine demethylation. Moreover, only Rpl3, actin and myosin are known to contain this modification, suggesting that a histidine demethylation activity might not be a priority for cells. If so, cells could produce 60S subunits with unmethylated Rpl3 by extracting methylated Rpl3 and replacing it with unmethylated protein. The chaperoned Rpl23, Rpl42 and Rpl43 are also methylated (Lee et al., 2002; Rossler et al., 2019; Ting et al., 2017; Yang et al., 2016). Mutation of the methylated residue in Rpl42 has no phenotype in rich medium, but confers resistance to high salt, glycerol and cold stress, as well as sensitivity to prolonged growth in stationary phase (Shirai et al., 2010). Thus, Rpl42 methylation is required to adapt to stationary growth, while under acute stress Rpl42 must be demethylated. Again, production of Rpl42-demethylated ribosomes might occur by Nap1-mediated extraction and subsequent replacement of Rpl42.

uS5/Rps2 is arginine-methylated in stationary phase (Ladror et al., 2014), and in fission yeast and higher eukaryotes forms a complex with the Tsr4 homolog Pdcd2 and the arginine methyltransferase Prmt3 (Bachand and Silver, 2004; Landry-Voyer et al., 2016; Lipson et al., 2010; Swiercz et al., 2007; Swiercz et al., 2005). Arginine methylation is expected to weaken Rps2 binding to ribosomes by reducing the net positive charge on the protein. Thus, arginine methylation could allow for Tsr4-mediated Rps2 extraction.

Finally, uL16/Rpl10 has UV-inducible isoforms in plants (Falcone Ferreyra et al., 2010), which could replace the generic isoform after its extraction by plant Sqt1.

These data show how exchange of a subset of ribosomal proteins could support a physiological stress response, and we therefore speculate that chaperone-mediated extraction could provide a mechanism for doing so.

### Rps26-deficient ribosomes, chaperones and cancer

Haploinsufficiency of Rps26 is a common cause of DBA (Doherty et al., 2010). Moreover, haploinsufficiency of Tsr2 is also linked to DBA (Gripp et al., 2014). Among the other most commonly associated genes are those for two other chaperoned proteins, Rpl5 and Rpl11 (Boria et al., 2010; Farrar et al., 2011). One of the phenotypic outcomes of DBA is the high incidence for specific forms of cancer including blood, breast, ovary, and colon cancers (Vlachos et al., 2012). Conversely, spontaneously formed cancers often lose the stoichiometry of their ribosomal proteins, especially when p53 is inactivated (Ajore et al., 2017; Guimaraes and Zavolan, 2016; Kulkarni et al., 2017; Vlachos, 2017). Finally, the differential resistance and sensitivities to certain cellular stresses (Collins et al., 2018; Ferretti et al., 2018; Parker et al., 2019), suggest that ribosomes lacking certain proteins perturb protein homeostasis, thereby supporting the development or progression of cancer.

In the absence of Tsr2, Rps26-deficient ribosomes accumulate (Schutz et al., 2014; Schutz et al., 2018). We show here that Tsr2 can also extract Rps26 from mature 40S ribosomes in the presence of high Na^+^ concentrations or high pH. Thus, Tsr2 is required to produce both Rps26-containing and deficient ribosomes. Intriguingly, cancer cells display elevated intracellular Na^+^ concentrations (Amara and Tiriveedhi, 2017) as well as pH values (White et al., 2017) and should therefore accumulate Rps26-deficient ribosomes, as long as they contain Tsr2, and perhaps even more so when Tsr2 is overexpressed. Thus, while in healthy cells Tsr2 deficiency leads to the accumulation of Rps26-deficient ribosomes, cancer cells, which display high intracellular salt and pH concentrations, require Tsr2 to accumulate Rps26-deficient ribosomes. Given the role for Rps26-deficient ribosomes in cancer, these data suggest that while Tsr2 deficiency might promote cancer development, Tsr2 is required for cancer maintenance. These seemingly contradictory roles for Tsr2 in cancer are consistent with observations in the cancer genome databases: The previously suggested role of Tsr2 in Rps26 incorporation indicates that there should be two routes to generate Rps26-deficient ribosomes: low expression of Rps26 or Tsr2. Indeed, database analysis provides evidence for both of these routes, most notably in over 40% of adenoid cystic breast carcinoma, or about 75% of gastric cancer samples, respectively (**Figure S5B**, (Cerami et al., 2012; Gao et al., 2013)). The function of Tsr2 in extraction of Rps26 discovered here also suggests a third route to Rps26-deficient ribosomes, the overexpression of Tsr2. Indeed, ~ 8% of prostate cancer samples in the database have *amplified* Tsr2, either with no changes in Rps26, or with the corresponding changes occurring in only a small subset of the samples, which could reflect upregulation of ribosome biogenesis (**Figure S5C**, (Cerami et al., 2012; Gao et al., 2013)). Finally, nearly 90 mutations in Tsr2, both within and outside of its Rps26 binding pocket have been described in cancer cells (**Figure S5D**, (Cerami et al., 2012; Gao et al., 2013)). Together, these analyses of the cancer genome datasets further support the role of Tsr2 in ribosomal Rps26-homeostasis for healthy cell growth via both its role in Rps26 incorporation and extraction. Similar superficially conflicting roles have been described for the other RP-chaperones: expression of BCCIP mRNA, encoding the human homolog for the Rpl23 chaperone Bcp1, is reduced in some cancer samples, but increased in others, and both can be linked to poor outcomes (Huang et al., 2013; Liu et al., 2009; Liu et al., 2013; Meng et al., 2003; Meng et al., 2007). Thus, it has been suggested that while haploinsufficiency can promote development of tumors, the protein is required for tumor progression (Droz-Rosario et al., 2017). GRWD1, the human homolog for the Rpl3 chaperone Rrb1, is titrated by the circular RNA circCDYL, the most highly expressed known circRNA, which displays numerous GRWD1 binding sites others (Okholm et al., 2020). Low circCDYNL expression, which should lead to high levels of free GRWD1, is associated with bladder cancer progression (Okholm et al., 2020), and consistently, GRWD1 overexpression also promotes tumorigenesis (Sugimoto et al., 2015). While GRWD1 is suggested to play cellular roles beyond chaperoning Rpl3 (Iouk et al., 2001; Pausch et al., 2015), we speculate that the tumorigenic effect from GRWD1 overexpression or circCDYL repression might arise from GRWD1-dependent Rpl3 extraction from ribosomes, leading to the production of Rpl3-deficient ribosomes.

### Cells may limit the amount of Rps26-deficient ribosomes via feedback cycles

The considerations above suggest the importance of ribosome homeostasis to ensure that populations of ribosomes lacking individual proteins are tightly regulated. Consistently, deletion of Rps26 is lethal in rich media, suggesting that Rps26-deficient ribosomes alone cannot support the translational needs of the cell. Moreover, haploinsufficiency of Rps26 leads to DBA in humans (Doherty et al., 2010), demonstrating the need to ensure that not too many Rps26-deficient ribosomes accumulate and that these do not persist for too long. How then do cells ensure the production of limited amounts of Rps26-deficient ribosomes? The mRNAs enriched on Rps26-deficient ribosomes include the Mg^2+^ transporter Alr2 and the Na^+^/H^+^ antiporter Nha1 (Ferretti et al., 2017). Thus, increased amounts of Rps26-deficient ribosomes should lead to increased production of Alr2 and Nha1, thereby increasing the cellular concentration of free Mg^2+^ and H^+^, while lowering intracellular concentrations of Na^+^. We have shown above that decreased Na^+^ promotes the reincorporation of Rps26 from Tsr2. Thus, this might be a self-regulating system, thereby ensuring that Rps26-deficient ribosomes do not dominate the ribosome population, even at high pH or Na^+^.

## STAR Methods

### Strains and plasmids

*S. cerevisiae* strains used in this study were either purchased from the GE Dharmacon Yeast Knockout Collection or constructed using standard methods (Longtine et al., 1998) and are listed in Table S1. Plasmids are listed in Table S2.

### Protein purification

Rps26, Tsr2 and Tsr2^DWI^ were expressed in *E. coli* Rosetta2 (DE3) cells (Novagen) as TEV-cleavable His6-MBP fusion proteins. Cells were grown at 37 °C in LB medium supplemented with antibiotics. At OD600 0.4 protein expression was induced with 1 mM IPTG for 16 h at 18 °C. Proteins were purified using Ni-NTA resin (Qiagen) according to the manufacturer’s instructions. Eluted proteins were pooled and dialyzed overnight at 4 °C into 50 mM Tris (pH 7.4), 100 mM NaCl, and 1 mM DTT with tobacco etch virus (TEV) protease. Tsr2 was further purified by MonoQ and Superdex 75 (GE) chromatography. For purification of the Rps26·Tsr2 complex, the His-MBP tag was removed by a second round of purification with Ni-NTA resin, and the Rps26·Tsr2 complex was further purified by Superdex 75 with complex buffer (50 mM Tris, pH 7.4, 500 mM NaCl). Concentrated proteins were stored at −80°C.

### Isolation of Rps3 TAP-tagged pre-made ribosomes

Cells were grown to mid-log phase in YPD to produce ribosomes with Rps3-TAP. After half of the cells were collected as unstressed cells, the other half were further incubated in YPGal with 1 M of NaCl for 4 hours in the presence of 0.2 µg/ml dox. After collecting the stress-treated cells, TAP-purification was performed (Ferretti et al., 2017), and the samples analyzed using Western blot. To test the repair of Rps26-deficient ribosomes, cells were further grown in YPGal for 1 hour after stress in the presence of dox.

### Isolation of Rps26 bound to Tsr2 after stress

Tsr2-TAP cells with pKK30528 (Gal:Rps26-HA) were grown to mid-log phase in YPGal media before expression of Rps26-HA was repressed by growth in YPD for 2 hours. Cells were then split into two pools and grown for 4h in YPGal with or without 1 M of NaCl. After collecting the cells, TAP-purification was performed as above, except 500mM NaCl was added to the wash buffer to remove ribosomes bound to Tsr2.

### In vitro Rps26 release assay

Pelleting release assays were performed as previously described (Khoshnevis et al., 2016). Briefly, 4 µM purified recombinant Tsr2 or Tsr2^DWI^ were mixed with 40 nM 40S subunits purified as previously described (Acker et al., 2007), incubated for 15 min at RT and 10 min on ice in binding buffer (20 mM HEPEs, pH 7.3, 2.5mM MgOAc, 0.1 mg/ml Heparin, 2 mM DTT and 0.5 µl RNasin [NEB]) containing different concentrations of salt or magnesium. Samples were layered onto a 400 μl sucrose cushion (ribosome binding buffer + 20% sucrose (w/v)) and spun for 2.5 h at 400,000 × g at 4°C in a TLA100.1 rotor (Beckman). Supernatants were precipitated using trichloroacetic acid and resuspended in the same volume as pellets. For pH dependent release assays, Bis-Tris propane was used to adjust the pH to the indicated value.

### qRT-PCR

Transcriptional repression of dox-repressible Rps3-TAP was assessed by inoculating YPGal media (with or without 0.2 μg/ml dox) with a preculture grown in YPD to mid-log phase. Cells were harvested at different time points, total RNA isolated by hot-phenol extraction, reverse transcribed and analyzed by qRT-PCR as described previously (Ferretti et al., 2017). qPCR primers are listed in **Table S3**.

### Antibodies

To detect TAP-tagged (Rps3 or Tsr2) or HA-tagged proteins (Rps26 variants), anti-TEV cleavage site from Invitrogen (PA1-119) or anti-HA antibody from Abcam (ab18181) or Sigma (ab1603) were used, respectively. For Rps10 and Rps26 detection, antibodies were raised by New England Peptide. The Tsr2/Rps26 antibody was a gift from V. Panse.

## Acknowledgements

We thank members of the Karbstein laboratory for discussion and comments on the manuscript. This work was supported by NIH grants R01-GM117093, R01-GM086451, R35-GM136323, and HHMI Faculty Scholar grant 55108536 to K.K., and an F32 fellowship to Y.Y. (F32-GM139302).

## Notes

### Competing Interest Statement

The authors have declared no competing interest.

## ReferencesUncategorized References

Ajore, R., Raiser, D., McConkey, M., Joud, M., Boidol, B., Mar, B., Saksena, G., Weinstock, D.M., Armstrong, S., Ellis, S.R., et al. (2017). Deletion of ribosomal protein genes is a common vulnerability in human cancer, especially in concert with TP53 mutations. EMBO Mol Med 9, 498–507.

Amara, S., and Tiriveedhi, V. (2017). Inflammatory role of high salt level in tumor microenvironment (Review). Int J Oncol 50, 1477–1481.

An, H., Ordureau, A., Korner, M., Paulo, J.A., and Harper, J.W. (2020). Systematic quantitative analysis of ribosome inventory during nutrient stress. Nature 583, 303–309.

Armistead, J., and Triggs-Raine, B. (2014). Diverse diseases from a ubiquitous process: the ribosomopathy paradox. FEBS Lett 588, 1491–1500.

Bachand, F., and Silver, P.A. (2004). PRMT3 is a ribosomal protein methyltransferase that affects the cellular levels of ribosomal subunits. EMBO J 23, 2641–2650.

Ben-Shem, A., Garreau de Loubresse, N., Melnikov, S., Jenner, L., Yusupova, G., and Yusupov, M. (2011). The structure of the eukaryotic ribosome at 3.0 A resolution. Science 334, 1524–1529.

Black, J.J., Musalgaonkar, S., and Johnson, A.W. (2019). Tsr4 Is a Cytoplasmic Chaperone for the Ribosomal Protein Rps2 in Saccharomyces cerevisiae. Mol Cell Biol 39.

Bolze, A., Mahlaoui, N., Byun, M., Turner, B., Trede, N., Ellis, S.R., Abhyankar, A., Itan, Y., Patin, E., Brebner, S., et al. (2013). Ribosomal protein SA haploinsufficiency in humans with isolated congenital asplenia. Science 340, 976–978.

Boria, I., Garelli, E., Gazda, H.T., Aspesi, A., Quarello, P., Pavesi, E., Ferrante, D., Meerpohl, J.J., Kartal, M., Da Costa, L., et al. (2010). The ribosomal basis of Diamond-Blackfan Anemia: mutation and database update. Hum Mutat 31, 1269–1279.

Brar, G.A., and Weissman, J.S. (2015). Ribosome profiling reveals the what, when, where and how of protein synthesis. Nat Rev Mol Cell Biol 16, 651–664.

Calvino, F.R., Kharde, S., Ori, A., Hendricks, A., Wild, K., Kressler, D., Bange, G., Hurt, E., Beck, M., and Sinning, I. (2015). Symportin 1 chaperones 5S RNP assembly during ribosome biogenesis by occupying an essential rRNA-binding site. Nat Commun 6, 6510.

Cerami, E., Gao, J., Dogrusoz, U., Gross, B.E., Sumer, S.O., Aksoy, B.A., Jacobsen, A., Byrne, C.J., Heuer, M.L., Larsson, E., et al. (2012). The cBio cancer genomics portal: an open platform for exploring multidimensional cancer genomics data. Cancer Discov 2, 401–404.

Chen, S.S., Sperling, E., Silverman, J.M., Davis, J.H., and Williamson, J.R. (2012). Measuring the dynamics of E. coli ribosome biogenesis using pulse-labeling and quantitative mass spectrometry. Mol Biosyst 8, 3325–3334.

Cheng, Z., Mugler, C.F., Keskin, A., Hodapp, S., Chan, L.Y., Weis, K., Mertins, P., Regev, A., Jovanovic, M., and Brar, G.A. (2019). Small and Large Ribosomal Subunit Deficiencies Lead to Distinct Gene Expression Signatures that Reflect Cellular Growth Rate. Mol Cell 73, 36–47 e10.

Cole, S.E., LaRiviere, F.J., Merrikh, C.N., and Moore, M.J. (2009). A convergence of rRNA and mRNA quality control pathways revealed by mechanistic analysis of nonfunctional rRNA decay. Mol Cell 34, 440–450.

Collins, J.C., Ghalei, H., Doherty, J.R., Huang, H., Culver, R.N., and Karbstein, K. (2018). Ribosome biogenesis factor Ltv1 chaperones the assembly of the small subunit head. J Cell Biol 217, 4141–4154.

Doherty, L., Sheen, M.R., Vlachos, A., Choesmel, V., O’Donohue, M.F., Clinton, C., Schneider, H.E., Sieff, C.A., Newburger, P.E., Ball, S.E., et al. (2010). Ribosomal protein genes RPS10 and RPS26 are commonly mutated in Diamond-Blackfan anemia. Am J Hum Genet 86, 222–228.

Droz-Rosario, R., Lu, H., Liu, J., Liu, N.A., Ganesan, S., Xia, B., Haffty, B.G., and Shen, Z. (2017). Roles of BCCIP deficiency in mammary tumorigenesis. Breast Cancer Res 19, 115.

Emmott, E., Jovanovic, M., and Slavov, N. (2019). Ribosome Stoichiometry: From Form to Function. Trends Biochem Sci 44, 95–109.

Falcone Ferreyra, M.L., Pezza, A., Biarc, J., Burlingame, A.L., and Casati, P. (2010). Plant L10 ribosomal proteins have different roles during development and translation under ultraviolet-B stress. Plant Physiol 153, 1878–1894.

Farrar, J.E., Vlachos, A., Atsidaftos, E., Carlson-Donohoe, H., Markello, T.C., Arceci, R.J., Ellis, S.R., Lipton, J.M., and Bodine, D.M. (2011). Ribosomal protein gene deletions in Diamond-Blackfan anemia. Blood 118, 6943–6951.

Ferrando, A., Kron, S.J., Rios, G., Fink, G.R., and Serrano, R. (1995). Regulation of cation transport in Saccharomyces cerevisiae by the salt tolerance gene HAL3. Mol Cell Biol 15, 5470–5481.

Ferretti, M.B., Barre, J.L., and Karbstein, K. (2018). Translational Reprogramming Provides a Blueprint for Cellular Adaptation. Cell Chem Biol 25, 1372–1379 e1373.

Ferretti, M.B., Ghalei, H., Ward, E.A., Potts, E.L., and Karbstein, K. (2017). Rps26 directs mRNA-specific translation by recognition of Kozak sequence elements. Nat Struct Mol Biol 24, 700–707.

Ferretti, M.B., and Karbstein, K. (2019). Does functional specialization of ribosomes really exist? RNA 25, 521–538.

Fortier, S., MacRae, T., Bilodeau, M., Sargeant, T., and Sauvageau, G. (2015). Haploinsufficiency screen highlights two distinct groups of ribosomal protein genes essential for embryonic stem cell fate. Proc Natl Acad Sci U S A 112, 2127–2132.

Gao, J., Aksoy, B.A., Dogrusoz, U., Dresdner, G., Gross, B., Sumer, S.O., Sun, Y., Jacobsen, A., Sinha, R., Larsson, E., et al. (2013). Integrative analysis of complex cancer genomics and clinical profiles using the cBioPortal. Sci Signal 6, pl1.

Gasch, A.P., Spellman, P.T., Kao, C.M., Carmel-Harel, O., Eisen, M.B., Storz, G., Botstein, D., and Brown, P.O. (2000). Genomic expression programs in the response of yeast cells to environmental changes. Mol Biol Cell 11, 4241–4257.

Genuth, N.R., and Barna, M. (2018a). The Discovery of Ribosome Heterogeneity and Its Implications for Gene Regulation and Organismal Life. Mol Cell 71, 364–374.

Genuth, N.R., and Barna, M. (2018b). Heterogeneity and specialized functions of translation machinery: from genes to organisms. Nat Rev Genet 19, 431–452.

Ghalei, H., Trepreau, J., Collins, J.C., Bhaskaran, H., Strunk, B.S., and Karbstein, K. (2017). The ATPase Fap7 Tests the Ability to Carry Out Translocation-like Conformational Changes and Releases Dim1 during 40S Ribosome Maturation. Mol Cell 68, 1155.

Gripp, K.W., Curry, C., Olney, A.H., Sandoval, C., Fisher, J., Chong, J.X., Genomics, U.W.C.f.M., Pilchman, L., Sahraoui, R., Stabley, D.L., et al. (2014). Diamond-Blackfan anemia with mandibulofacial dystostosis is heterogeneous, including the novel DBA genes TSR2 and RPS28. Am J Med Genet A 164A, 2240–2249.

Guimaraes, J.C., and Zavolan, M. (2016). Patterns of ribosomal protein expression specify normal and malignant human cells. Genome Biol 17, 236.

Huang, H., Ghalei, H., and Karbstein, K. (2020). Quality control of 40S ribosome head assembly ensures scanning competence. J Cell Biol 219.

Huang, Y.Y., Dai, L., Gaines, D., Droz-Rosario, R., Lu, H., Liu, J., and Shen, Z. (2013). BCCIP suppresses tumor initiation but is required for tumor progression. Cancer Res 73, 7122–7133.

Iouk, T.L., Aitchison, J.D., Maguire, S., and Wozniak, R.W. (2001). Rrb1p, a yeast nuclear WD-repeat protein involved in the regulation of ribosome biosynthesis. Mol Cell Biol 21, 1260–1271.

Khoshnevis, S., Askenasy, I., Johnson, M.C., Dattolo, M.D., Young-Erdos, C.L., Stroupe, M.E., and Karbstein, K. (2016). The DEAD-box Protein Rok1 Orchestrates 40S and 60S Ribosome Assembly by Promoting the Release of Rrp5 from Pre-40S Ribosomes to Allow for 60S Maturation. PLoS Biol 14, e1002480.

Koch, B., Mitterer, V., Niederhauser, J., Stanborough, T., Murat, G., Rechberger, G., Bergler, H., Kressler, D., and Pertschy, B. (2012). Yar1 protects the ribosomal protein Rps3 from aggregation. J Biol Chem 287, 21806–21815.

Kondrashov, N., Pusic, A., Stumpf, C.R., Shimizu, K., Hsieh, A.C., Ishijima, J., Shiroishi, T., and Barna, M. (2011). Ribosome-mediated specificity in Hox mRNA translation and vertebrate tissue patterning. Cell 145, 383–397.

Koplin, A., Preissler, S., Ilina, Y., Koch, M., Scior, A., Erhardt, M., and Deuerling, E. (2010). A dual function for chaperones SSB-RAC and the NAC nascent polypeptide-associated complex on ribosomes. J Cell Biol 189, 57–68.

Kressler, D., Bange, G., Ogawa, Y., Stjepanovic, G., Bradatsch, B., Pratte, D., Amlacher, S., Strauss, D., Yoneda, Y., Katahira, J., et al. (2012). Synchronizing nuclear import of ribosomal proteins with ribosome assembly. Science 338, 666–671.

Kulkarni, S., Dolezal, J.M., Wang, H., Jackson, L., Lu, J., Frodey, B.P., Dosunmu-Ogunbi, A., Li, Y., Fromherz, M., Kang, A., et al. (2017). Ribosomopathy-like properties of murine and human cancers. PLoS One 12, e0182705.

Ladror, D.T., Frey, B.L., Scalf, M., Levenstein, M.E., Artymiuk, J.M., and Smith, L.M. (2014). Methylation of yeast ribosomal protein S2 is elevated during stationary phase growth conditions. Biochem Biophys Res Commun 445, 535–541.

Landry-Voyer, A.M., Bilodeau, S., Bergeron, D., Dionne, K.L., Port, S.A., Rouleau, C., Boisvert, F.M., Kehlenbach, R.H., and Bachand, F. (2016). Human PDCD2L Is an Export Substrate of CRM1 That Associates with 40S Ribosomal Subunit Precursors. Mol Cell Biol 36, 3019–3032.

LaRiviere, F.J., Cole, S.E., Ferullo, D.J., and Moore, M.J. (2006). A late-acting quality control process for mature eukaryotic rRNAs. Mol Cell 24, 619–626.

Larsson, K., Bohl, F., Sjostrom, I., Akhtar, N., Strand, D., Mechler, B.M., Grabowski, R., and Adler, L. (1998). The Saccharomyces cerevisiae SOP1 and SOP2 genes, which act in cation homeostasis, can be functionally substituted by the Drosophila lethal(2)giant larvae tumor suppressor gene. J Biol Chem 273, 33610–33618.

Lebaron, S., Schneider, C., van Nues, R.W., Swiatkowska, A., Walsh, D., Bottcher, B., Granneman, S., Watkins, N.J., and Tollervey, D. (2012). Proofreading of pre-40S ribosome maturation by a translation initiation factor and 60S subunits. Nat Struct Mol Biol 19, 744–753.

Lee, S.W., Berger, S.J., Martinovic, S., Pasa-Tolic, L., Anderson, G.A., Shen, Y., Zhao, R., and Smith, R.D. (2002). Direct mass spectrometric analysis of intact proteins of the yeast large ribosomal subunit using capillary LC/FTICR. Proc Natl Acad Sci U S A 99, 5942–5947.

Lilleorg, S., Reier, K., Pulk, A., Liiv, A., Tammsalu, T., Peil, L., Cate, J.H.D., and Remme, J. (2019). Bacterial ribosome heterogeneity: Changes in ribosomal protein composition during transition into stationary growth phase. Biochimie 156, 169–180.

Lipson, R.S., Webb, K.J., and Clarke, S.G. (2010). Rmt1 catalyzes zinc-finger independent arginine methylation of ribosomal protein Rps2 in Saccharomyces cerevisiae. Biochem Biophys Res Commun 391, 1658–1662.

Liu, J., Lu, H., Ohgaki, H., Merlo, A., and Shen, Z. (2009). Alterations of BCCIP, a BRCA2 interacting protein, in astrocytomas. BMC Cancer 9, 268.

Liu, X., Cao, L., Ni, J., Liu, N., Zhao, X., Wang, Y., Zhu, L., Wang, L., Wang, J., Yue, Y., et al. (2013). Differential BCCIP gene expression in primary human ovarian cancer, renal cell carcinoma and colorectal cancer tissues. Int J Oncol 43, 1925–1934.

Longtine, M.S., McKenzie, A., 3rd, Demarini, D.J., Shah, N.G., Wach, A., Brachat, A., Philippsen, P., and Pringle, J.R. (1998). Additional modules for versatile and economical PCR-based gene deletion and modification in Saccharomyces cerevisiae. Yeast 14, 953–961.

Loveland, A.B., Bah, E., Madireddy, R., Zhang, Y., Brilot, A.F., Grigorieff, N., and Korostelev, A.A. (2016). Ribosome*RelA structures reveal the mechanism of stringent response activation. Elife 5.

MacGilvray, M.E., Shishkova, E., Chasman, D., Place, M., Gitter, A., Coon, J.J., and Gasch, A.P. (2018). Network inference reveals novel connections in pathways regulating growth and defense in the yeast salt response. PLoS Comput Biol 13, e1006088.

Mathis, A.D., Naylor, B.C., Carson, R.H., Evans, E., Harwell, J., Knecht, J., Hexem, E., Peelor, F.F., 3rd, Miller, B.F., Hamilton, K.L., et al. (2017). Mechanisms of In Vivo Ribosome Maintenance Change in Response to Nutrient Signals. Mol Cell Proteomics 16, 243–254.

Meng, X., Liu, J., and Shen, Z. (2003). Genomic structure of the human BCCIP gene and its expression in cancer. Gene 302, 139–146.

Meng, X., Yue, J., Liu, Z., and Shen, Z. (2007). Abrogation of the transactivation activity of p53 by BCCIP down-regulation. J Biol Chem 282, 1570–1576.

Mitterer, V., Gantenbein, N., Birner-Gruenberger, R., Murat, G., Bergler, H., Kressler, D., and Pertschy, B. (2016). Nuclear import of dimerized ribosomal protein Rps3 in complex with its chaperone Yar1. Sci Rep 6, 36714.

Okholm, T.L.H., Sathe, S., Park, S.S., Kamstrup, A.B., Rasmussen, A.M., Shankar, A., Chua, Z.M., Fristrup, N., Nielsen, M.M., Vang, S., et al. (2020). Transcriptome-wide profiles of circular RNA and RNA-binding protein interactions reveal effects on circular RNA biogenesis and cancer pathway expression. Genome Med 12, 112.

Orij, R., Postmus, J., Ter Beek, A., Brul, S., and Smits, G.J. (2009). In vivo measurement of cytosolic and mitochondrial pH using a pH-sensitive GFP derivative in Saccharomyces cerevisiae reveals a relation between intracellular pH and growth. Microbiology (Reading) 155, 268–278.

Parker, M.D., Collins, J.C., Korona, B., Ghalei, H., and Karbstein, K. (2019). A kinase-dependent checkpoint prevents escape of immature ribosomes into the translating pool. PLoS Biol 17, e3000329.

Pausch, P., Singh, U., Ahmed, Y.L., Pillet, B., Murat, G., Altegoer, F., Stier, G., Thoms, M., Hurt, E., Sinning, I., et al. (2015). Co-translational capturing of nascent ribosomal proteins by their dedicated chaperones. Nat Commun 6, 7494.

Peng, W.T., Robinson, M.D., Mnaimneh, S., Krogan, N.J., Cagney, G., Morris, Q., Davierwala, A.P., Grigull, J., Yang, X., Zhang, W., et al. (2003). A panoramic view of yeast noncoding RNA processing. Cell 113, 919–933.

Pillet, B., Garcia-Gomez, J.J., Pausch, P., Falquet, L., Bange, G., de la Cruz, J., and Kressler, D. (2015). The Dedicated Chaperone Acl4 Escorts Ribosomal Protein Rpl4 to Its Nuclear Pre-60S Assembly Site. PLoS Genet 11, e1005565.

Pulk, A., Liiv, A., Peil, L., Maivali, U., Nierhaus, K., and Remme, J. (2010). Ribosome reactivation by replacement of damaged proteins. Mol Microbiol 75, 801–814.

Robertson, W.R., Dowsett, S.J., and Hardy, S.J. (1977). Exchange of ribosomal proteins among the ribosomes of Escherichia coli. Mol Gen Genet 157, 205–214.

Rossler, I., Embacher, J., Pillet, B., Murat, G., Liesinger, L., Hafner, J., Unterluggauer, J.J., Birner-Gruenberger, R., Kressler, D., and Pertschy, B. (2019). Tsr4 and Nap1, two novel members of the ribosomal protein chaperOME. Nucleic Acids Res 47, 6984–7002.

Sauert, M., Temmel, H., and Moll, I. (2015). Heterogeneity of the translational machinery: Variations on a common theme. Biochimie 114, 39–47.

Schutz, S., Fischer, U., Altvater, M., Nerurkar, P., Pena, C., Gerber, M., Chang, Y., Caesar, S., Schubert, O.T., Schlenstedt, G., et al. (2014). A RanGTP-independent mechanism allows ribosomal protein nuclear import for ribosome assembly. Elife 3, e03473.

Schutz, S., Michel, E., Damberger, F.F., Oplova, M., Pena, C., Leitner, A., Aebersold, R., Allain, F.H., and Panse, V.G. (2018). Molecular basis for disassembly of an importin:ribosomal protein complex by the escortin Tsr2. Nat Commun 9, 3669.

Shi, Z., Fujii, K., Kovary, K.M., Genuth, N.R., Rost, H.L., Teruel, M.N., and Barna, M. (2017). Heterogeneous Ribosomes Preferentially Translate Distinct Subpools of mRNAs Genome-wide. Mol Cell 67, 71–83 e77.

Shirai, A., Sadaie, M., Shinmyozu, K., and Nakayama, J. (2010). Methylation of ribosomal protein L42 regulates ribosomal function and stress-adapted cell growth. J Biol Chem 285, 22448–22460.

Slavov, N., Semrau, S., Airoldi, E., Budnik, B., and van Oudenaarden, A. (2015). Differential Stoichiometry among Core Ribosomal Proteins. Cell Rep 13, 865–873.

Stelter, P., Huber, F.M., Kunze, R., Flemming, D., Hoelz, A., and Hurt, E. (2015). Coordinated Ribosomal L4 Protein Assembly into the Pre-Ribosome Is Regulated by Its Eukaryote-Specific Extension. Mol Cell 58, 854–862.

Strunk, B.S., Novak, M.N., Young, C.L., and Karbstein, K. (2012). A translation-like cycle is a quality control checkpoint for maturing 40S ribosome subunits. Cell 150, 111–121.

Subramanian, A.R., and van Duin, J. (1977). Exchange of individual ribosomal proteins between ribosomes as studied by heavy isotope-transfer experiments. Mol Gen Genet 158, 1–9.

Sugimoto, N., Maehara, K., Yoshida, K., Yasukouchi, S., Osano, S., Watanabe, S., Aizawa, M., Yugawa, T., Kiyono, T., Kurumizaka, H., et al. (2015). Cdt1-binding protein GRWD1 is a novel histone-binding protein that facilitates MCM loading through its influence on chromatin architecture. Nucleic Acids Res 43, 5898–5911.

Sulima, S.O., Patchett, S., Advani, V.M., De Keersmaecker, K., Johnson, A.W., and Dinman, J.D. (2014). Bypass of the pre-60S ribosomal quality control as a pathway to oncogenesis. Proc Natl Acad Sci U S A 111, 5640–5645.

Swiercz, R., Cheng, D., Kim, D., and Bedford, M.T. (2007). Ribosomal protein rpS2 is hypomethylated in PRMT3-deficient mice. J Biol Chem 282, 16917–16923.

Swiercz, R., Person, M.D., and Bedford, M.T. (2005). Ribosomal protein S2 is a substrate for mammalian PRMT3 (protein arginine methyltransferase 3). Biochem J 386, 85–91.

Ting, Y.H., Lu, T.J., Johnson, A.W., Shie, J.T., Chen, B.R., Kumar, S.S., and Lo, K.Y. (2017). Bcp1 Is the Nuclear Chaperone of Rpl23 in Saccharomyces cerevisiae. J Biol Chem 292, 585–596.

Vlachos, A. (2017). Acquired ribosomopathies in leukemia and solid tumors. Hematology Am Soc Hematol Educ Program 2017, 716–719.

Vlachos, A., Rosenberg, P.S., Atsidaftos, E., Alter, B.P., and Lipton, J.M. (2012). Incidence of neoplasia in Diamond Blackfan anemia: a report from the Diamond Blackfan Anemia Registry. Blood 119, 3815–3819.

Warner, J.R. (1999). The economics of ribosome biosynthesis in yeast. Trends Biochem Sci 24, 437–440.

Webb, K.J., Zurita-Lopez, C.I., Al-Hadid, Q., Laganowsky, A., Young, B.D., Lipson, R.S., Souda, P., Faull, K.F., Whitelegge, J.P., and Clarke, S.G. (2010). A novel 3-methylhistidine modification of yeast ribosomal protein Rpl3 is dependent upon the YIL110W methyltransferase. J Biol Chem 285, 37598–37606.

West, M., Hedges, J.B., Chen, A., and Johnson, A.W. (2005). Defining the order in which Nmd3p and Rpl10p load onto nascent 60S ribosomal subunits. Mol Cell Biol 25, 3802–3813.

White, K.A., Grillo-Hill, B.K., and Barber, D.L. (2017). Cancer cell behaviors mediated by dysregulated pH dynamics at a glance. J Cell Sci 130, 663–669.

Yang, Y.T., Ting, Y.H., Liang, K.J., and Lo, K.Y. (2016). The Roles of Puf6 and Loc1 in 60S Biogenesis Are Interdependent, and Both Are Required for Efficient Accommodation of Rpl43. J Biol Chem 291, 19312–19323.

