## Supplemental for "The chaperone Tsr2 regulates Rps26 release and reincorporation from mature ribosomes to enable a reversible, ribosome-mediated response to stress"

This file contains

5 Supplemental Figures
3 Supplemental Tables
Supplemental References

### FigureS1

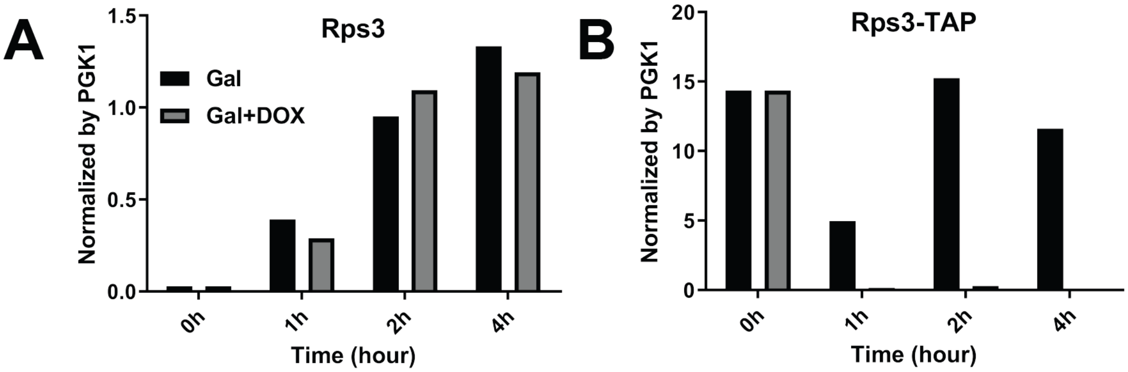


**Figure S1: Rps3 and Rps3-TAP mRNA induction and repression upon galactose and doxycycline (dox) addition.** Rps3 (A) and Rps3-TAP (B) mRNA levels normalized to PGK1 mRNA as measured by qRT-PCR. Cells grown in glucose media were switched to galactose media in mid-log phase with or without doxycycline. qPCR primers were designed to differentiate between Rps3 and Rps3-Tap (**Table S3**).

FigureS2

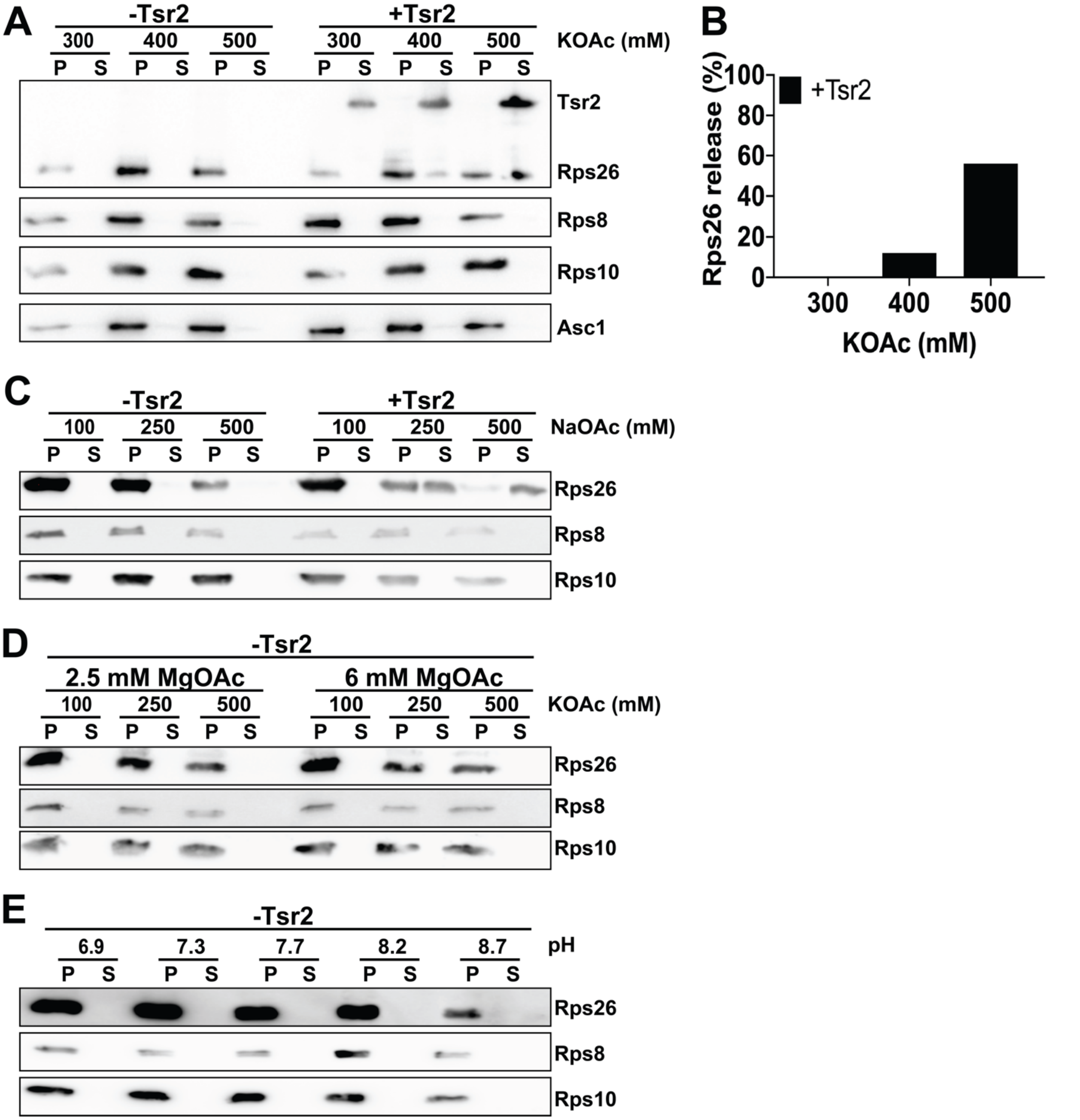


**Figure S2: Release of Rps26 by Tsr2 under increased salt.** (A) Release of Rps26, Rps8, Rps10 and Asc1 was measured in presence and absence of Tsr2 in different potassium concentrations by western blot. (B) Quantification of Rps26 in panel (A). (C) Tsr2 is required for $Na^+$ dependent release of Rps26. Release of Rps26, Rps8 and Rps10 was measured in presence and absence of Tsr2 at different $Na^+$ values by western blot. (D) Tsr2 is required for $Mg^{2+}$

dependent release of Rps26. Release of Rps26, Rps8 and Rps10 was measured in absence of Tsr2 at different $Mg^{2+}$ values by western blot. (E) Tsr2 is required for pH dependent release of Rps26. Release of Rps26, Rps8 and Rps10 was measured in absence of Tsr2 at different pH values by western blot.

### FigureS3

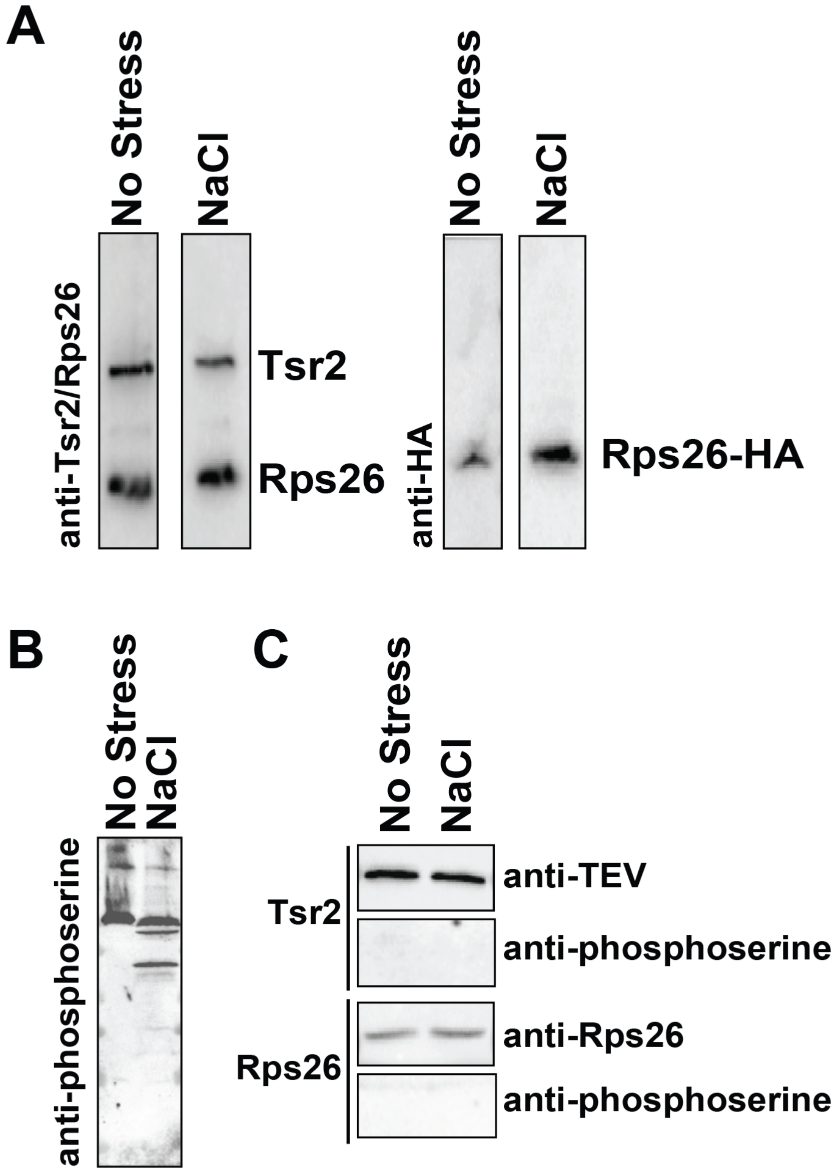


**Figure S3: No post-translational modification observed in Rps26 and Tsr2 after stress.** (A) Western blot of Rps26 and Rps26-HA from newly made and pre-existing ribosomes, respectively, co-isolated with TAP-tagged Tsr2 from cells that were or were not treated with 1 M NaCl. Elution samples were detected using primary antibodies of anti-Tsr2/Rps26 or anti-HA. (B) Western blot of total cell lysate of cells treated with or without 1 M NaCl. (C) Western blot of Tsr2-TAP purified from lysate in panel (B). Anti-phosphoserine antibody from sigma (AB1603) was used to detect phosphorylated proteins in panel (B) and (C).

### FigureS4

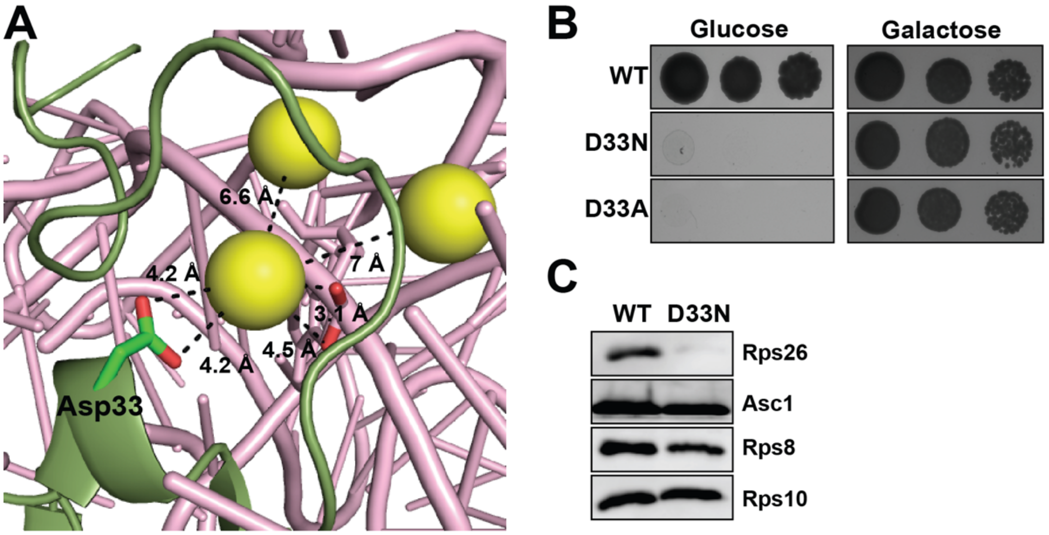


**Figure S4. Effect of Asp33 mutation in Rps26**. (A) Aspartate 33 in Rps26 binds a metal ion, displayed in yellow spheres (PDB 4V88). (B) Mutation of Asp33 mutation to asparagine or alanine leads to growth defects in yeast cells. (C) Occupancy of Rps26 in 40S subunits purified from cells expressing either Rps26_WT or Rps26_D33N.

### FigureS5

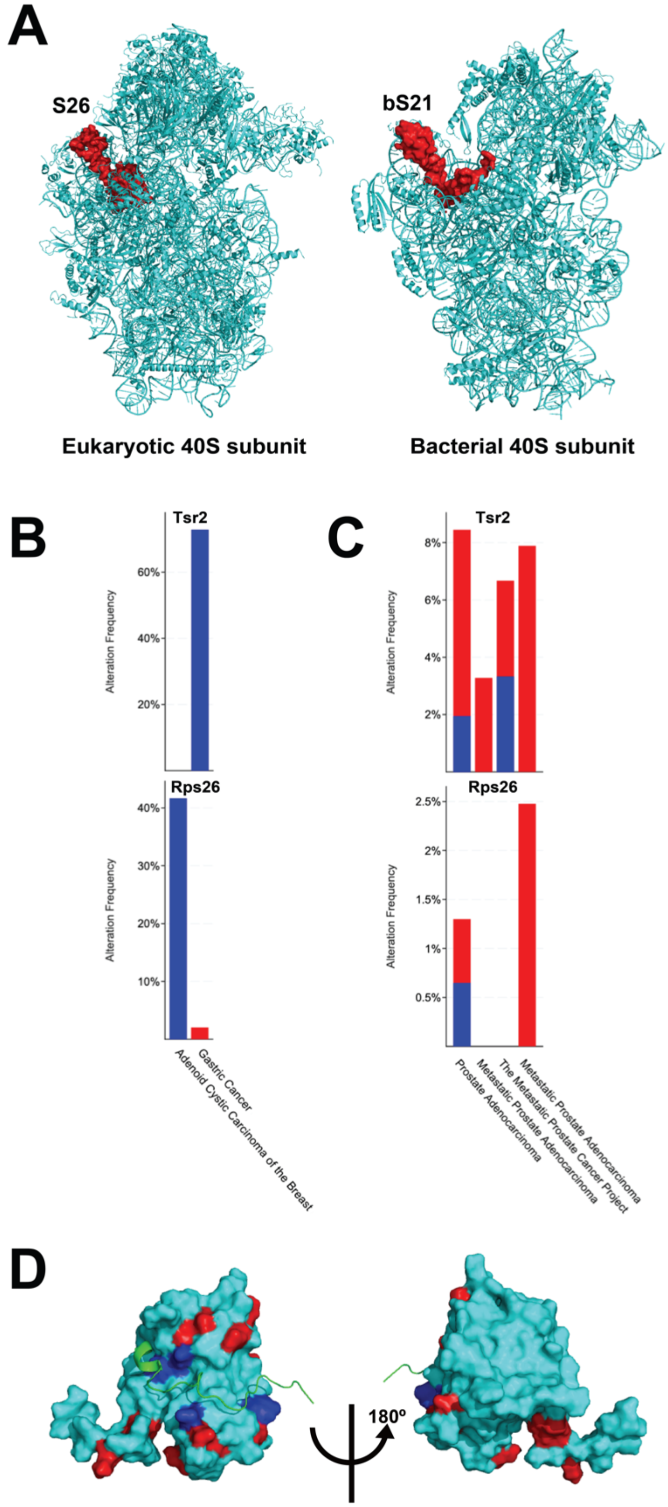
A
S26
bS21
Eukaryotic 40S subunit
Bacterial 40S subunit
B
Tsr2
Rps26
Alteration Frequency
60%
40%
20%
30%
10%
Adenoid Cystic Carcinoma of the Breast
Gastric Cancer
C
Tsr2
Rps26
Alteration Frequency
8%
6%
4%
2%
2.5%
1.5%
1%
0.5%
Prostate Adenocarcinoma
Metastatic Prostate Adenocarcinoma
The Metastatic Prostate Cancer Project
Metastatic Prostate Adenocarcinoma
D
180º

**Figure S5.** (A) Position of Rps26 in the eukaryotic 40S subunit (PDB 4V88) and bS21 in the bacterial 30S subunit (PDB 7K00). (B and C) cBio Portal analysis of genomic amplification (red) or deep deletion (blue) of Tsr2 or Rps26 in selected cancer cell lines. (D) Mutations in Tsr2 (cyan) described in cancer cells are marked with blue (Rps26 binding pocket) or red (outside binding pocket). The Rps26 tail is colored green. PDB 6G04.

**Table S1: Yeast strains used in this work**

| Strain | Description | Background | Genotype | Reference |
|---|---|---|---|---|
| YKK200 | WT | BY4741 | *MATα his3Δ1 leu2Δ0 met15Δ0 ura3Δ0* | GE Dharmacon |
| YKK491 | Gal::Rps26 | BY4741 | *MATα NatMX6::pGAL1-Rps26A Rps26B::KanMX6 his3Δ1 leu2Δ0 met15Δ0 ura3Δ0* | (Ferretti et al., 2017) |
| YKK493 | Gal::Rps3 | BY4741 | *MATα KanMX6::pGAL1-Rps3 his3Δ1 leu2Δ0 met15Δ0 ura3Δ0* | (Ferretti et al., 2017) |
| YKK856 | Tsr2-TAP | BY4741 | *MATα NatMX6::Tsr2-TAP his3Δ1 leu2Δ0 met15Δ0 ura3Δ0* | GE Dharmacon |
| YKK1109 | Gal-Tsr2 | BY4741 | *MATα NatMX6::pGAL1-Tsr2 his3Δ1 leu2Δ0 met15Δ0 ura3Δ0* | (Schutz et al., 2014) |

**Table S2: Plasmids used in this work**

| Plasmid | Description | Backbone | Reference |
|---|---|---|---|
| pKK3558 | TEF::Rps26A | pRS416 | (Ferretti et al., 2017) |
| pKK30832 | TEF::Rps26A_D33N | pRS416 | This work |
| pKK30831 | TEF::Rps26A_D33A | pRS416 | This work |
| pKK30528 | Gal::Rps26A-HA | pRS426 | This work |
| pKK30592 | TEF::Rps26A-HA | pRS416 | This work |
| pKK30562 | TEF::Rps26A$^{1\text{-}99}$-HA | pRS416 | This work |
| pKK30012 | TET:: Rps3-TAP | pCM189 | This work |

**Table S3: qPCR primers used in this work**

| Gene | Type | Sequence |
|---|---|---|
| Rps3 | FW | GCTGTCACCATCATTGAACC |
| | RV | GCACTAGAATAGAAGAAATTATTG |
| Rps3-TAP | FW | GCTGTCACCATCATTGAACC |
| | RV | CAAGTGCCCCGGAGGATGAG |
| PGK1 | FW | GCTGCTTTGCCAACCATCAA |
| | RV | GGCTTCAACTTCTGGACCGA |
